# RAPDOR: Using Jensen-Shannon Distance for the computational analysis of complex proteomics datasets

**DOI:** 10.1101/2024.09.30.615781

**Authors:** Luisa Hemm, Dominik Rabsch, Halie R. Ropp, Viktoria Reimann, Philip Gerth, Jürgen Bartel, Manuel Brenes-Álvarez, Sandra Maaß, Dörte Becher, Wolfgang R. Hess, Rolf Backofen

## Abstract

The computational analysis of large proteomics datasets, such as those from gradient profiling or spatially resolved proteomics, is often as crucial as the experimental design. We present RAPDOR, a tool for intuitive analyzing and visualizing such datasets, based on the Jensen-Shannon distance and subsequent analysis of similarities between replicates, applied to three datasets. First, we examined the in-gradient distribution profiles of protein complexes with or without RNase treatment (GradR) to identify the set of RNA-binding proteins (RBPs) in the cyanobacterium *Synechocystis* sp. PCC 6803. RBPs play pivotal regulatory and structural roles; although numerous RBPs have been identified, the complete set is unknown for any species. RAPDOR identified 80 potential RBPs, including ribosomal proteins, likely RNA-modifying enzymes, and several proteins not previously associated with RNA binding. High-ranking putative RBPs, such as the universal stress protein Sll1388, or the translation inhibitor LrtA/RaiA, were predicted by RAPDOR but not the TriPepSVM algorithm, indicating uncharacterized RBP domains. These data are available online at https://synecho-rapdor.biologie.uni-freiburg.de, providing a comprehensive resource for RNase-sensitive protein complexes in cyanobacteria. We then show by reanalyzing existing datasets, that RAPDOR is effective in examining the intracellular redistribution of proteins under stress conditions. RAPDOR is a generic, non-parametric tool for the intuitive and versatile analysis of highly complex data sets such as the study of protein distributions using fractionation protocols.

## Introduction

RNA-binding proteins (RBPs) are crucial components of ribonucleoprotein complexes, including ribosomes, the signal recognition particle, and CRISPR-Cas complexes and play vital roles in all domains of life. The eukaryotic RBP database lists more than 6,300 ortholog groups with more than 315,000 individual RBPs across 162 eukaryotic species^1^. Several thousand RBPs have more recently been identified in mammals, including many metabolic enzymes that are also binding to RNA^2,3^. RBPs play crucial roles in various regulatory pathways and are involved in the regulation of alternative splicing^4^, in neural cell maturation in mammal cells^5^, cancer and epigenetic mechanisms^6^, and development in plants^7^.

In prokaryotes, RBPs play significant roles in the post-transcriptional regulation of gene expression. Previously described regulatory RBPs in gram-negative bacteria include Hfq^8,9^, ProQ^10–12^, CsrA^13^ and KhpA/B^14^. Many new RBPs have been discovered in recent years also in other groups of bacteria, such as in gram-positive *Streptococcus pneumoniae*^15^. Several RBPs were described also for different Archaea^16^. However, the complete set of RBPs has not yet been identified for any organism.

Consequently, experimental, and computational methods have been developed to identify putative RBPs. *In silico* methods mostly rely on amino acid strings (k-mers) as features to classify the RNA binding nature of a protein. Prominent examples for such prediction tools based on machine learning are RBPPred^17^, its successor Deep-RBPPred^18^, and TriPepSVM^19^. TriPepSVM was trained on k-mer frequencies of known RBPs from different organisms. However, it can identify RBPs also in cross-species predictions.

High throughput experimental methods for identifying candidate RBPs include fractionation approaches, which are based on separating the cell lysate by density gradient ultracentrifugation or size exclusion chromatography and extracting fractions based on differences in molecular mass or buoyant density. In the Grad-Seq ultracentrifugation approach, the proteome and transcriptome composition of each fraction is determined. Overlapping protein/transcript occurrences indicate potential RBP-RNA interactions. The first application of this method identified the major RNA chaperone ProQ in *Salmonella*^20^. Grad-Seq has been used since to identify novel RBP candidates in different types of bacteria, including *Clostridioides difficile*, *Enterococcus* species, *Fusobacterium nucleatum*, and the cyanobacterium *Synechocystis* sp. PCC 6803 (from here *Synechocystis* 6803)^14,15,21–25^.

To obtain a higher resolution of the captured complexome, the glycerol or sucrose ultracentrifugation gradient can be replaced with size exclusion chromatography^26^. However, the co-occurrence of a particular RNA and a particular protein is not necessarily indicating their interaction. To address this issue, two further protocols were established. R-DeeP was developed using the HeLa S3 cell line^27^, while GradR was developed in *Salmonella enterica*^28^. Both methods are based on gradient fractionation of whole cell lysates, similar to Grad-Seq. But in contrast to Grad-Seq, two gradients are prepared in parallel. One gradient is loaded with cell lysate that was treated with RNase beforehand, while the other gradient serves as a control and is loaded with untreated cell lysate. After ultracentrifugation and fractionation, the protein contents of all fractions are measured using mass spectrometry. RBPs can be identified by a shift in the fractions of the RNase-treated gradient, as the additional mass of the RNA is removed upon digestion. The R-DeeP study identified 1,784 RNA-dependent proteins in HeLa S3 cells. Of these, 537 proteins lacked a previous link to RNA^27^. In *Salmonella*, the RBP FopA was identified using this method^28^.

Since these types of experiments produce large and complex datasets, their computational analysis is just as important as the experimental design. While analysis pipelines have been published along with these experiments, their flexibility regarding the number of fractions and replicates is often limited. For example, R-DeeP fits Gaussian models to the curve representing the mean mass spec profile from the three replicates for each condition. To ensure as many peaks as maxima are found in the profile, the location, value and standard deviation estimate for each maximum found was provided for the Gaussian model. In a second step, Gaussian models were fitted to each replicate. A Student’s t test was then used to assess the p-value (FDR-corrected) for difference between the Gaussian fits of control and treatment, indicating shifts that are associated with RNA dependencies^27^. However, the protein abundances throughout the different fractions do not necessarily follow Gaussian distributions and the R script used to analyze the experiment was fixed to 25 fractions and three replicates per group. Adjusting this script for a different number of fractions requires considerable manual effort.

In another approach, hierarchical clustering was employed to analyze GradR data and discover new RBPs clustering with known ones^28^. Although in this way the FopA protein was discovered as a new member of the family of FinO/ProQ-like RBPs^28^, this methodology lacks a straightforward way to account for experimental and biological variance by using replicates.

To overcome existing limitations, we developed a tool based on the Jensen-Shannon Distance (*JSD*) and the analysis of similarities (ANOSIM), called **R**apid **A**NOSIM using **P**robability **D**istance for estimati**O**n of **R**edistribution (**RAPDOR**). RAPDOR can be used to analyze any distribution of proteins over a fractionation analysis with two conditions (e.g. RNase treated vs. control). Since RAPDOR is independent of the fractionation approach, it can handle data resulting from ultracentrifugation as well as size exclusion chromatography.

As a direct application, we used RAPDOR for the analysis of protein complexes after gradient profiling with or without RNase treatment (GradR) in the model cyanobacterium *Synechocystis* 6803. Cyanobacteria differ from most non-photosynthetic bacteria by the presence of extensive intracellular photosynthetic membrane systems, the thylakoids. But it is the presence of these membrane systems that likely triggered a particular set of RRM domain-containing RBPs to get involved in the intracellular transport and localization of certain mRNAs^29^. Moreover, there is extensive evidence for the presence of post-transcriptional regulation in cyanobacteria^30,31,31–35^. However, information on regulatory RBPs and RNA chaperones in cyanobacteria is limited. Homologs of enterobacterial RNA chaperones, such as CsrA, ProQ, FinO, or Hfq are missing in cyanobacteria, or do not bind RNA^36^. For *Synechocystis* 6803, previous Grad-Seq analysis yielded a small number of potential novel RBPs^25^. One of these proteins, the YlxR homologue Ssr1238, was recently verified as involved in tRNA maturation^37^.

Here, we provide a list of potential RBPs in *Synechocystis* 6803 that were identified by RAPDOR using newly generated GradR data and computationally predicted by a custom version of the TriPepSVM algorithm. In addition we applied RAPDOR to existing spatial proteomics datasets from HeLa cells^38^ and *Synechocystis* 6803^39^. Both datasets investigate the redistribution of proteins among cellular compartments under various stress conditions. While the original publications employed an approach tailored for the dataset, using parametric tests to identify these distribution shifts, our findings demonstrate that the generic and non-parametric RAPDOR workflow produces similar results with fewer statistical assumptions, thus being more robust and less prone to outliers in different application scenarios. RAPDOR as a flexible-to-use tool is available as a pypi package with its source code and its documentation on https://domonik.github.io/RAPDOR/. The code used to analyze the data is available as a snakemake workflow on Github (https://github.com/domonik/synRDPMSpec). The GradR data is available online in a static version of RADPOR at https://synecho-rapdor.biologie.uni-freiburg.de, providing a comprehensive resource for the identification of RNase-sensitive protein complexes in *Synechocystis* 6803.

## Results

### GradR analysis of an unicellular cyanobacterium

Triplicates of *Synechocystis* 6803 cultures were grown under moderate light conditions (50 µmol m^−2^ s^−1^) in BG11 medium and cell lysates were prepared when an OD_750_ of 0.9 was reached. The cleared cell lysates were split into two parts of identical volumes. One part of each lysate was RNase-treated, while the second was mock-treated for control (**Fig. 1A**). Then, all lysates were subjected to GradR ultracentrifugation, but using β-D-maltoside (DDM) as a soft membrane solubilizer and sucrose gradients instead of glycerol. Gradients showed a distinct color profile after centrifugation, relating to the specific pigment-containing complexes of *Synechocystis* **(Fig. 1B, C** and **D**). In the untreated sample the RNA was distributed along the gradient with peaks in concentration in fractions 3-6 for short RNAs and fractions 16-20 for longer RNAs. Most RNA was in pellet fraction 20 (**Fig. 1B**). As a proof of concept, the distribution of some known RNA within the fractions is shown in **Fig. S1.** The RNase P RNA RnpB^41^ was detected in fractions 6-13, with a peak in fractions 9 and 10. A similar distribution was obtained for the transfer-messenger RNA (tmRNA) SsrA^42^, while the sRNA PmgR1^31^ was detected in fractions 5-12, with a peak in fractions 8 and 9 (**Fig. S1**).

**Fig. 1.**
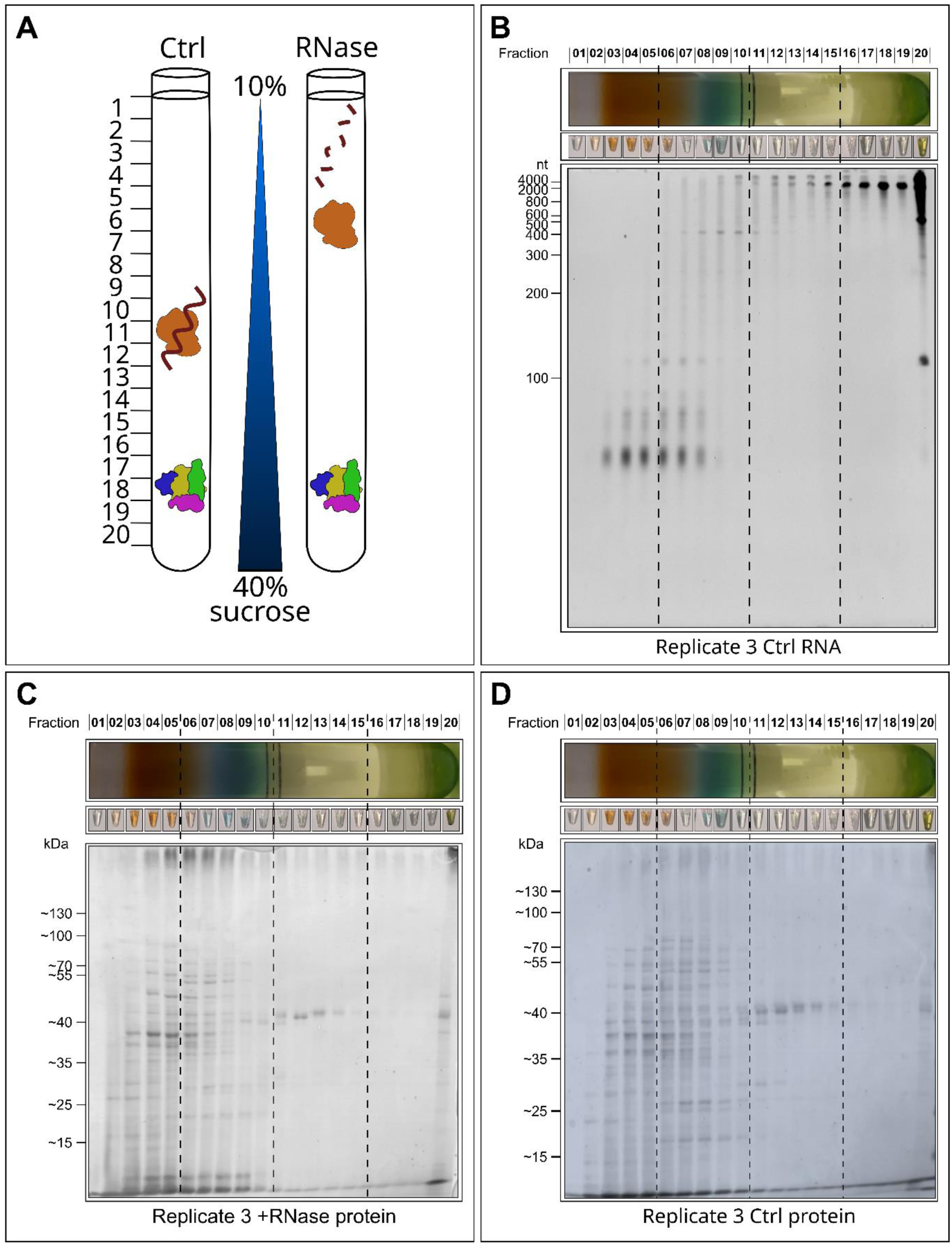
Experimental approach and fractionation of gradients. (**A**) Experimental set-up to identify RNA dependent proteins. **(B)** Separation of the extracted RNA of the untreated control sample replicate 3 on a 10% denaturing gel stained with ethidium bromide verifies RNA quality from the different fractions. **(C)** Protein distribution for the RNase-treated fractions. **(D)** Protein distribution for the fractions without RNase. In panels C and D, a 12% SDS-PAGE was loaded with 20 µL per fraction of replicate 3 and stained with InstantBlue Coomassie Protein Stain (Abcam).

The protein composition of each fraction was determined by mass spectrometry analysis, measuring 100 µL aliquots of each fraction. In total 1,134 proteins were detected, 6 were found in only one replicate, 62 in two replicates and the rest had unique peptides in all three replicates (**Supplementary Table S1**). Hence, ∼31% of the annotated 3.681 proteins in *Synechocystis* 6803 were identified. Besides the mass spectrometric analysis, the distribution of proteins was visualized by SDS-PAGE with subsequent Coomassie blue staining (**Fig. 1C and D**). In the gel, no difference between the treated and untreated samples was observed. Most proteins were present in fractions 3-8 and in the pellet fraction 20. An overview of fraction complexity and distribution is given in the **Supplementary Table S2**. Plotting the sedimentation of selected protein complexes to the respective fractions in comparison to the calculated molecular masses indicated a resolution limit of ∼50 kDa **(Fig. S2**). A regression analysis showed a linear relationship between the calculated molecular masses and their fraction distribution.

### A novel tool for the analysis of protein compartment/fraction distribution profiles

For the analysis of in-gradient distribution profiles, we developed a novel tool based on the *JSD*^43^, called RAPDOR. The analysis workflow of such profiles consists of basically three steps: 1) preprocessing, 2) detection of significantly differential profiles between the treated and untreated samples, and 3) prediction of the profile shift between conditions (**Fig. 2**). Concerning preprocessing, the input for the RAPDOR workflow is a csv file containing a row for each protein and a column for each fraction, and sample, respectively. Our tool further allows to use an averaging kernel of adjustable size to smooth the distributions along the fraction dimension. This can reduce the variance among replicates since experimental fractionation is usually not 100% reproducible. The smoothed intensities are then normalized to add up to 1. This results in a probability distribution function 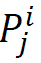 where *x* ∈ *X* is the fraction number, *i* marks the corresponding protein and *j* the sample. In preparation for the second step (detection of significantly different profiles), it calculates the mean distribution 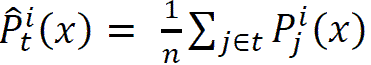 for the treated (*t* = +) and untreated (*t* = −) *n* replicates for each protein.

**FIG 2.**
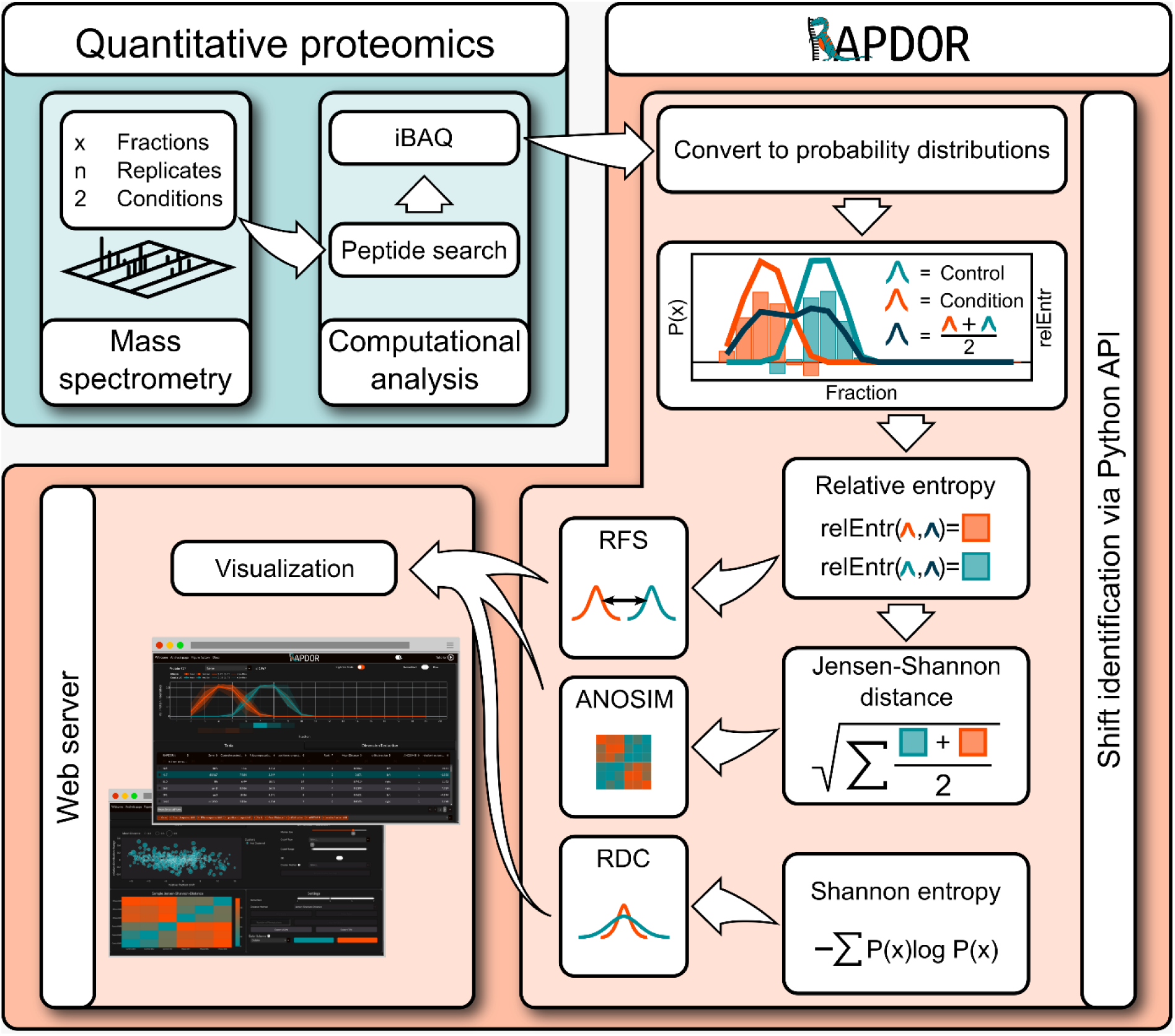
RAPDOR workflow. The RAPDOR tool converts analyzed mass spectrometry data with x fractions from n replicates and two conditions into probability distributions. It continues with calculation of the relative entropy (relEntr) and the Shannon entropy. It uses the Jensen-Shannon distance (*JSD*) between all pairwise samples to carry out an ANOSIM and evaluates the effect size via the *JSD* between the mean distributions of the conditions. Lastly, distribution changes are visualized in an interactive webserver displaying several parameters such as the relative position shift (RSD) and the relative distribution change (RDC). Depending on the number of replicates, the individual proteins can be either tested for redistribution using ANOSIM or ranked according to their assigned ANOSIM R values.

Concerning the second and third step, the RAPDOR workflow is the first tool that uses a non-parametric approach for detection of significantly different profiles. To the best of our knowledge, the only other approach to determine profile differences and shift directions is R-DeeP^27^. R-DeeP, however, uses a parametric approach by assuming a Gaussian model fit to the curve representing the distribution profile from the three replicates for each condition. In addition, it uses the parametric Student’s t test to evaluate the significance of peak shifts, which are found by the Gaussian fitting process. Especially the assumption of a Gaussian mixture model for the distribution profile is critical as there is only a small number of fractions, which leads to boundary effects that are poorly modelled by a Gaussian mixture model. Consequently, we find that R-DeeP has problems in analyzing proteins of large complexes, such as the ribosomal RNA-binding protein RpsQ, where the untreated condition usually has a large value in the last fraction, which is then distributed to nearby lower fractions in the treated condition (**Supplementary Fig. S3**).

For our non-parametric approach, we compare the profiles between treated and untreated samples by interpreting them as probability distributions 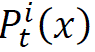 (*i* being the protein, *t* being the condition), as described before. A natural selection to determine the effect size is then a metric between probability distributions. Thus, we use the *JSD* as default to evaluate the effect size via the 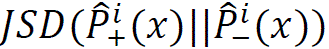 (Eq. (3)) of the mean distributions. Internally, the *JSD* uses the KL divergence (Eq. 2) of two distributions to their mixture distribution, resulting in a divergence measurement that is a metric and thus symmetric. Under the null hypothesis that the treated and untreated distributions are equal, the two distributions will be the same or at least very close, resulting in a small average KL-Divergence to the mean distribution. On the other hand, an increasing KL-Divergence to the mixture distribution gathers information against this null hypothesis. Additionally, the *JSD* is constrained within the range of 0 to 1 when using a logarithmic base of 2. A value of 0 shows that the two distributions do not overlap at all, while a value of 1 indicates that they are identical, thus enhancing the interpretability of the default effect size measure. Also, it is not negatively influenced by the high number of zero values measured for most proteins along the gradient in GradR. Thus, the *JSD* (i.e., our definition of effect size) between mean profiles per condition is part of the ranking provided by RAPDOR.

While the effect size as defined by the *JSD* is a valuable information for step 2 (detection of significantly differential profiles), it does not provide a rating of significance. For statistical significance, RAPDOR makes use of the replicate structure of the GradR experiments or other fractionation protocols, which have typically three replicates. Using *JSD* on all pairs to measure the effect size for a specific pair of replicates for the same protein, we generate a matrix of differences (or dissimilarity) for each protein. A popular non-parametric test on a matrix of dissimilarity was introduced with the ANOSIM R statistics (**An**alysis **o**f **sim**ilarities^44^). Here, the matrix of dissimilarity corresponds to a set of samples, each belonging to a single site. The idea of the ANOSIM R statistics is simply to evaluate the rank similarities within the same condition/site and between conditions/sites. The null hypothesis *H*_0_ is that the similarities between conditions are greater or equal to the similarities within a condition. The corresponding test statistics R is the difference between the average rank similarities of pairs of samples from the two different conditions (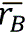) and the average rank similarities of pairs of samples from the same condition (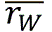, see Eq. 7(7) and methods).

As initially mentioned^44^, the “R statistic itself is a useful comparative measure of the degree of separation of sites” (in our case, conditions). This implies that we can use the calculated R, for each protein, to rank proteins according to their likelihood of binding RNA, providing a more sensitive ranking than the effect size on mean profiles. That allows, for example, a user of the RAPDOR workflow (**Fig. 2**) to look at the R statistics of some known RNA-binding proteins, and to investigate all proteins with an R higher than the known RNA-binding proteins.

Depending on the number of replicates, it is even possible to calculate a p-value. For this purpose, we implement the permutation test for *H*_0_ as described in the original ANOSIM paper (see also methods for details). The basic idea is to generate all possible permutations of test/untreated labels for the replicates of a specific protein, and then calculate the R value which provides an experimental distribution for the R values for this protein. However, for the typical number of n=3 replicates and k=2 conditions, there are only (2 ∗ 3)!⁄((3!)^2^ ∗ 2!)=10 (^44^, Eq. 8) possible distinct permutations, one being the original matrix. Thus, we cannot generate significant results with three replicates using the permutations for each protein individually. For that reason, RAPDOR allows to generate permutations for all proteins and conditions to determine a sampling distribution. For *i* proteins, this results in (*n* ∗ 2)! ∗ *i* many R values from distinct permutations for *a* dataset (**Supplementary Fig. S4**). Using the distribution of these value, RAPDOR enables the calculation of a p-value for each protein. However, when using a small number of replicates as typically used in these fields (e.g., three to four replicates), the statistical power will still be low after a correction for multiple testing and control of the false discovery rate. However, users can counteract this via either adjusting their significance level or using the R value ranking instead of p-values.

In the third step, we need an automatic way to evaluate the direction and the length of the shift. Here, RAPDOR computes, for each protein, the contribution of each position to the KL-divergence between the mean distributions 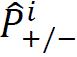 to the corresponding mixture distribution (Eq. (4)). Peaks can then be defined as the positions with maximal KL divergence contribution. However, plateau-like peak profiles would result in a noisy peak detection. For that reason, we apply a temperature-scaled soft-argmax function to determine the peak locations. This procedure highlights the expected position of the strongest shift *S*_*t*_ in the treated and untreated replicates. Via subtracting the two positions, the shift length is determined as a value called the relative fraction shift. A shift is called left if the subtraction results in a negative value and right if it is greater than zero. Noteworthy, this calculates a shift direction also for very similar distributions. Therefore, it is crucial to differentiate shifts based on the mean distance of their peaks and ANOSIM, consequently interpreting the direction of the shift only when it is clearly discernible.

Finally, we provide an additional visualization termed *bubble plot* (such as in **Fig. 5**) for investigating the experimental results further. Thus, one interesting information is whether, for a given protein, the mean distribution does have a similar Shannon entropy in both conditions or not. A similar entropy would e.g. occur when we have a clear peak that is only shifted in its location in the two conditions. The entropy would be different, though, if there is a clear peak in one condition, which is flattened in the other condition. Thus, the developed tool further evaluates whether a shift led to a broader or sharper distribution. Subtracting those two entropies yields a single number, which we call the relative distribution change. Hereby positive values mean that the protein had a much broader distribution after treatment. In contrast, a low negative value indicates that the protein accumulated in a single fraction and was before uniformly distributed along the gradient. In combination, the bubble plot displays the entropy difference (y-axis), the fraction shift and direction of the two calculated peaks (x-axis) together with the effect size (size of the bubble), yielding an excellent tool for a fast selection of candidate proteins. The overall workflow is described in **Fig. 2**.

### RAPDOR is well-suited to analyze GradR data

The ribosomal protein RplA was used as a control for the shifted position in gradient fractions after RNase digestion. RplA is a member of the uL1 ribosomal protein family and directly interacts with the 23S rRNA^45,46^, making it a suitable example for RNase-induced shifts. In the mass spectrometry data as well as in the experimental verification by Western blot analysis, RplA showed a very clear left shift from fraction 20 to fraction ∼4 **(Fig. 3A)**. This shift was picked up and visualized well in the RAPDOR analysis **(Fig. 3A)**. RAPDOR indicated for the small ribosomal subunit proteins a general shift to lower molecular weight fractions, whereas the large ribosomal unit proteins did not completely disintegrate upon RNase digestion, with a few proteins, such as RplA, showing strong shifts and others remaining in high molecular weight complexes **(Fig. 3B)**. This behavior was also visible in the histograms. Most of the proteins belonging to the small subunit showed a low effect size shift that was reproducible among the replicates as indicated by the distribution of the mean *JSDs* and the high ANOSIM R values. In contrast, the distribution of R values for proteins from the large subunit resembled the incomplete disintegration as only some proteins showed larger values, but most were centered around 0. When sorting the dataset according to a decreasing ANOSIM R and a decreasing *JSD*, the median rank of proteins from the small ribosomal subunit was much smaller than the median rank of proteins from the large ribosomal subunit **(Fig. 3C)**. No shift was observed for the purely protein-based photosystem I and II complexes indicating that their in-gradient distributions were not influenced by RNA-binding **(Fig. 3B)**. We conclude that the experiment as well as the RAPDOR algorithm worked well for identifying known RNA interaction partners.

**FIG 3.**
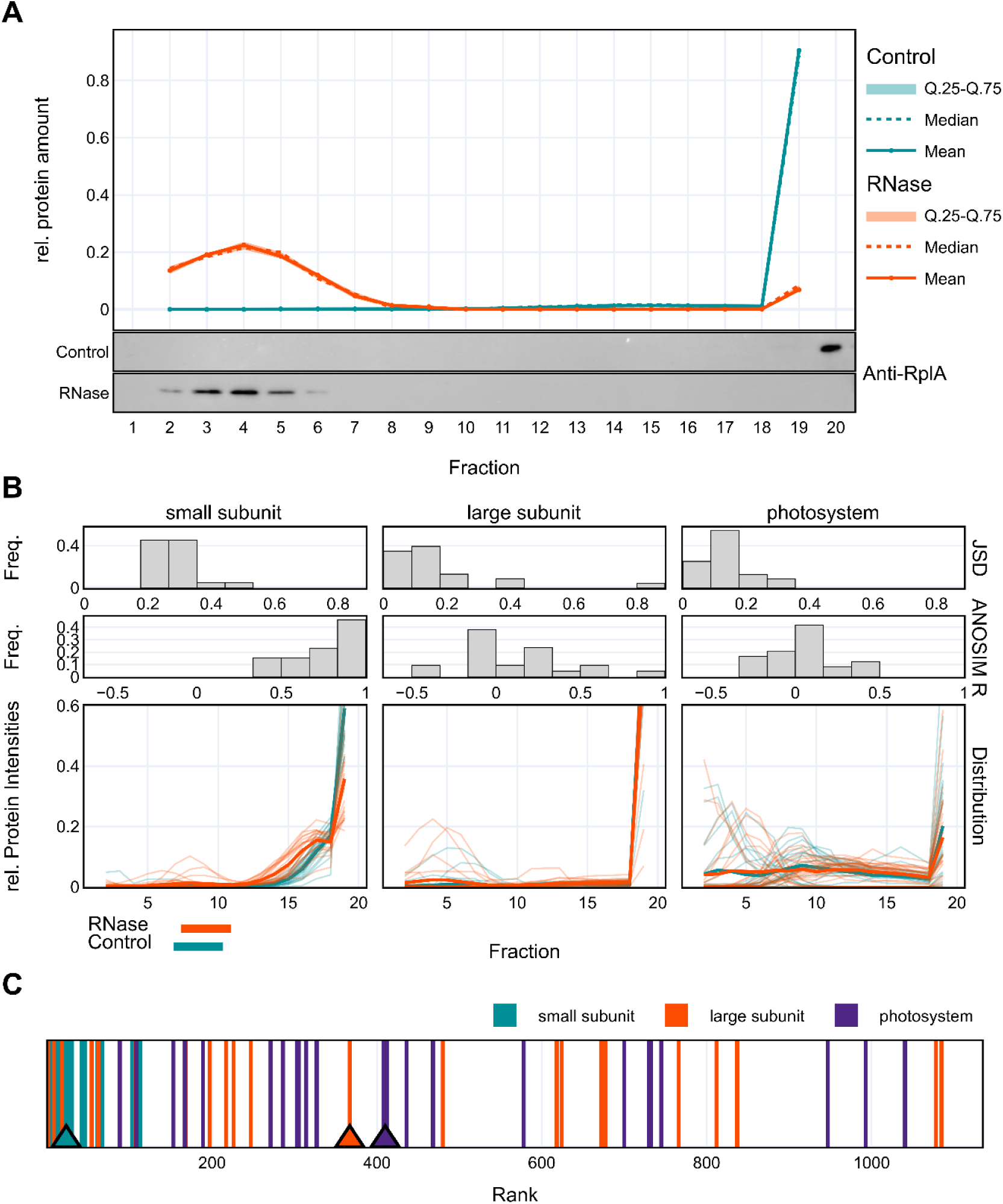
The shift of ribosomal proteins in the RNase-treated fractions shows proof of principle. (**A**) Shift of large subunit protein RplA was detected by western blot of replicate 1 samples and mass spectrometry data analyzed by RAPDOR (all replicates). **(B)** Almost all proteins of the ribosomal small subunit show a leftward shift supported by a high ANOSIM R and a *JSD* > 0.2. The large ribosomal subunit disassembles with few proteins showing a strong shift to lower molecular weight fractions, such as RplA. Proteins of the photosystems do not shift. **(C)** Plot of ranked RAPDOR median values for the three protein groups from panel B (triangles). The median for small subunit ribosomal proteins is at 23, while the medians of the other two groups were at much larger ranks (smaller values indicate higher probability of RNA binding).

### Comparison between analysis tools for GradR datasets

We compared proteins with an ANOSIM R ≥ 0.5 computed by RAPDOR to those showing a significant change (p-value ≤ 0.05) between the peaks observed in the RNase-treated samples compared to the controls using an adapted R-DeeP script^27^. Since the original R-DeeP script only supports experimental setups containing 25 fractions, we modified the code to work with 20 fractions whilst keeping other parameters unchanged. We decided for the ANOSIM cutoff via manually inspecting the R values of proteins known to bind RNA. In addition, we considered the distribution of ANOSIM R’s from all proteins generated using every possible permutation of the treatment labels (**Supplementary Fig. S4**). Although the number of proteins within these thresholds (ANOSIM R value ≥ 0.5, p-value ≤ 0.05) was limited and their selection might result in a relatively high false discovery rate, we opted for this procedure as it is still sensitive enough to identify promising candidates for further investigation. In addition, we considered the TripPepSVM-generated list of 306 candidate RBPs with a score ≥0.25 (**Supplementary Table S3**. For further benchmarking we checked the numbers of ribosomal proteins that were detected by the different approaches. TripPepSVM classified 47 of the 52 ribosomal proteins as RBPs (**Fig. 4**). By mass spectrometry, 52 ribosomal proteins were identified. Based on the experimental evidence, RAPDOR detected 24 ribosomal proteins as RBP, while 28 were below our ANOSIM R threshold of 0.5. In contrast, R-DeeP identified shifts only for 4 ribosomal proteins. In the RAPDOR analysis, 9 of the 24 ribosomal proteins exhibited the highest achievable ANOSIM R value of 1 and a *JSD* exceeding 0.2 (**Supplementary Table S4**).

**FIG 4.**
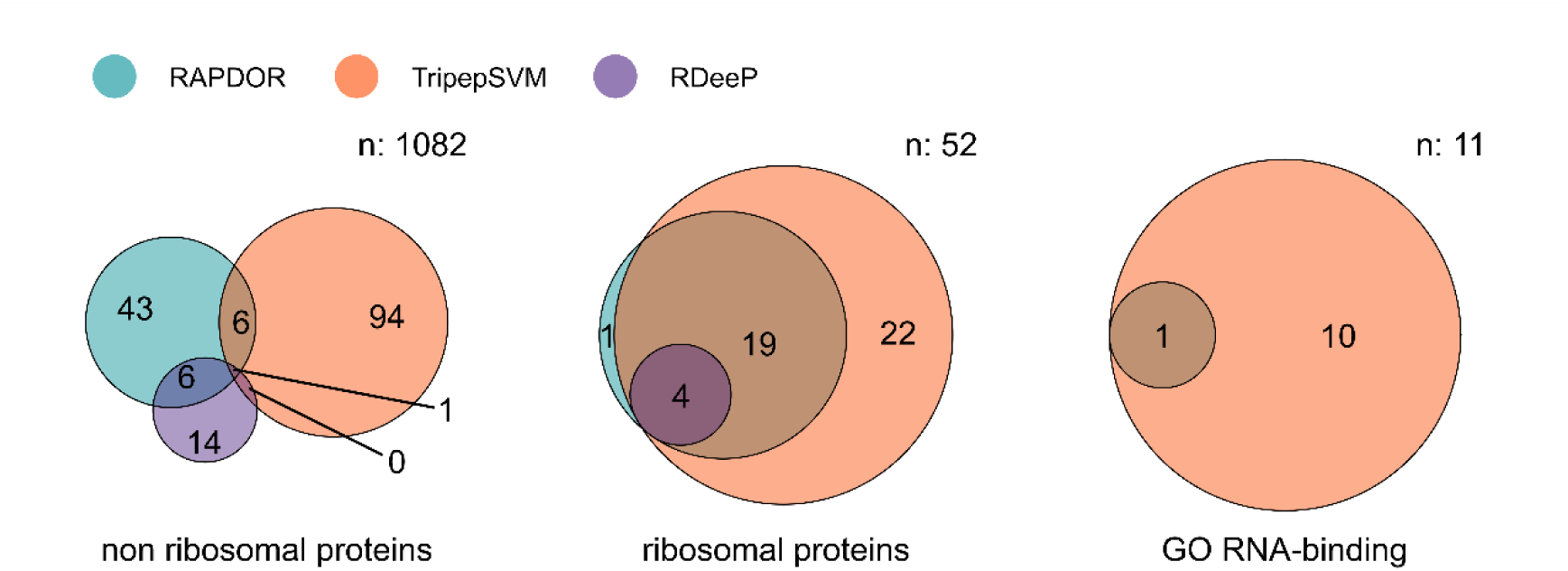
RNA-dependent proteins identified via the different approaches. The respective total numbers of proteins identified by mass spectrometry are indicated as n values. The first Venn diagram shows all identified RNA-dependent proteins excluding ribosomal proteins. The number of ribosomal proteins identified as RNA-dependent via the different approaches is displayed in the middle. The right panel summarizes all identified proteins with the RNA-binding GO annotation that were not ribosomal proteins.

Besides the ribosomal proteins, 11 other proteins detected by mass spectrometry, harbor the GO Term for RNA binding in the *Synechocystis* 6803 Uniprot annotation (from a total of 37 annotated RBPs). They were all included in the SVM training data and thus classified by TriPepSVM as RBPs. RAPDOR found one of these annotated RBPs, the RRM-domain containing protein RbpA on rank 33, and none were detected by R-DeeP. Both pipelines used to analyze the experimental data showed a low memory consumption of approximately 150 Mb (**Supplementary Fig. S5A**). However, RAPDOR outperformed the R-DeeP pipeline regarding runtime (**Supplementary Fig. S5A**). Additionally, it demonstrated superior speed compared to R-DeeP running with three replicates, even when incorporating p-value calculations on a simulated dataset featuring nine replicates per group.

### Candidates for proteins interacting with RNA from GradR data

Proteins with a high ANOSIM R value or an SVM prediction as RBP represent promising candidates for further research. Therefore, we compiled a table of non-ribosomal proteins falling into these categories using the following approach. After the RAPDOR analysis, the results were filtered for an ANOSIM R ≥0.5 and after removing ribosomal proteins from this list, 54 proteins remained for detailed analysis **(Supplementary Table S5)**. From these we selected 20 proteins for more detailed analysis based on additional criteria **(Table 1)**. These criteria were the behavior of homologues in GradR experiments in another cyanobacterium, *Nostoc* sp. PCC 7120^47^, homologies to other known RBPs or a promising SVM score as potential RBPs. Two proteins in this list are known RBPs, namely Ffh, the apoprotein of the signal recognition particle^48^ and the RRM domain protein RbpA/Rbp1^49^. These two proteins were highly ranked also by their SVM score (on ranks 14 and 17, **Table 1**), which is likely a direct effect of training the TriPepSVM algorithm with known RBPs.

**Table 1.** Selected RNA-dependent candidates based on RAPDOR analysis and manual filtering. After removing ribosomal proteins, 20 proteins with a promising rank in the RAPDOR analysis are given together with their SVM prediction of RNA binding (**Supplementary Table S3**). The gene name or locus tag is given, followed by the ANOSIM R value (ANO), the mean JSD distance (MDS), the relative shift in the number of fractions, the absolute rank assigned by RAPDOR and SVM, comments and annotation. n/a indicates an SVM score below threshold.

| Protein | ANO | MDS | Shift | Rank | SVM | Comments and annotation |
| --- | --- | --- | --- | --- | --- | --- |
| SII1967 | 1 | 0.87 | -3.92 | 2 | 282 | RNA methyltransferase RmID, TRAM domain |
| Ffh | 1 | 0.74 | +11.92 | 3 | 19 | Signal recognition particle protein |
| *SII1388 | 1 | 0.13 | -12 | 15 | n/a | Universal stress protein |
| Slr0782 | 0.89 | 0.33 | -3.21 | 22 | n/a | L-amino acid dehydrogenase |
| SII0921 | 0.83 | 0.54 | -5.18 | 28 | n/a | NarL subfamily, HTH LuxR-type domain |
| RaiA/LrtA | 0.81 | 0.35 | -12.74 | 29 | n/a | Ribosome hibernation promotion factor (Hpf) |
| QueF | 0.81 | 0.18 | -3.26 | 31 | n/a | 7-cyano-7-deazaguanine reductase Slr0711 |
| Slr0147 | 0.78 | 0.4 | -2.92 | 32 | n/a | 4-vinyl reductase |
| RbpA | 0.78 | 0.26 | -5.07 | 33 | 16 | RNA-binding protein RbpA/Rbp1; RRM domain |
| Ssl3355 | 0.78 | 0.2 | -11.29 | 35 | n/a | preprotein translocase subunit SecE |
| Pgm | 0.78 | 0.16 | -4.94 | 36 | n/a | Phosphoglucomutase |
| Slr1143 | 0.75 | 0.64 | -2.92 | 37 | n/a | GGDEF domain, diguanylate synthase |
| *Slr2130 | 0.74 | 0.29 | -3.19 | 39 | n/a | AroB, 3-dehydroquinate synthase |
| SII7087 | 0.63 | 0.38 | -6 | 51 | n/a | CRISPR-Cas protein Cmr4 |
| SII0284 | 0.59 | 0.26 | +3 | 59 | 248 | Bipolar DNA helicase HerA |
| *Slr0678 | 0.58 | 0.58 | -0.76 | 63 | n/a | Biopolymer transporter ExbD |
| SII1961 | 0.55 | 0.37 | -8.23 | 64 | n/a | GntR/AqpZ family transcriptional regulator |
| RpoC1 | 0.52 | 0.23 | -0.31 | 78 | n/a | RNAP subunit gamma |
| ChII | 0.44 | 0.3 | -7.31 | 93 | 228 | Mg<sup>2+</sup> chelatase subunit ChII |
| CphA | 0.33 | 0.13 | -8.33 | 148 | 277 | Cyanophycin synthetase |

All proteins from the list are marked in **Fig. 5**, proteins that shifted to lower molecular weight fractions upon RNase digestion are on the left side of this plot, while proteins that shifted to higher molecular weight fractions are on the right side. Six of the selected candidates were located in the lower left quarter, around RbpA. Upon RNase digestion, these proteins shifted to lower molecular weight fractions and narrowed in distribution. Four of the selected proteins (Hpf, Sll1388, ChlI and Sll0921) were located in the upper left quarter, meaning that the distribution along the gradient broadened, but the respective proteins still shifted to the lower molecular weight fractions, two of these were also suggested by TriPepSVM (ChlI on rank 112, Hpf on rank 169, **Table 1**). Only one of the selected candidates showed a rightward shift and a narrowed distribution, located in the lower right quadrant. This protein is the signal recognition particle protein (Ffh), a proven RBP, which forms ribonucleoprotein complexes and serves as an important factor in protein translocation^48^. Upon RNase digestion, Ffh shifted from fractions 6-8 to the pellet fraction, showing a strong right shift **(Fig. 6)**. This behavior indicates the loss of solubility upon RNase digestion.

**FIG 5.**
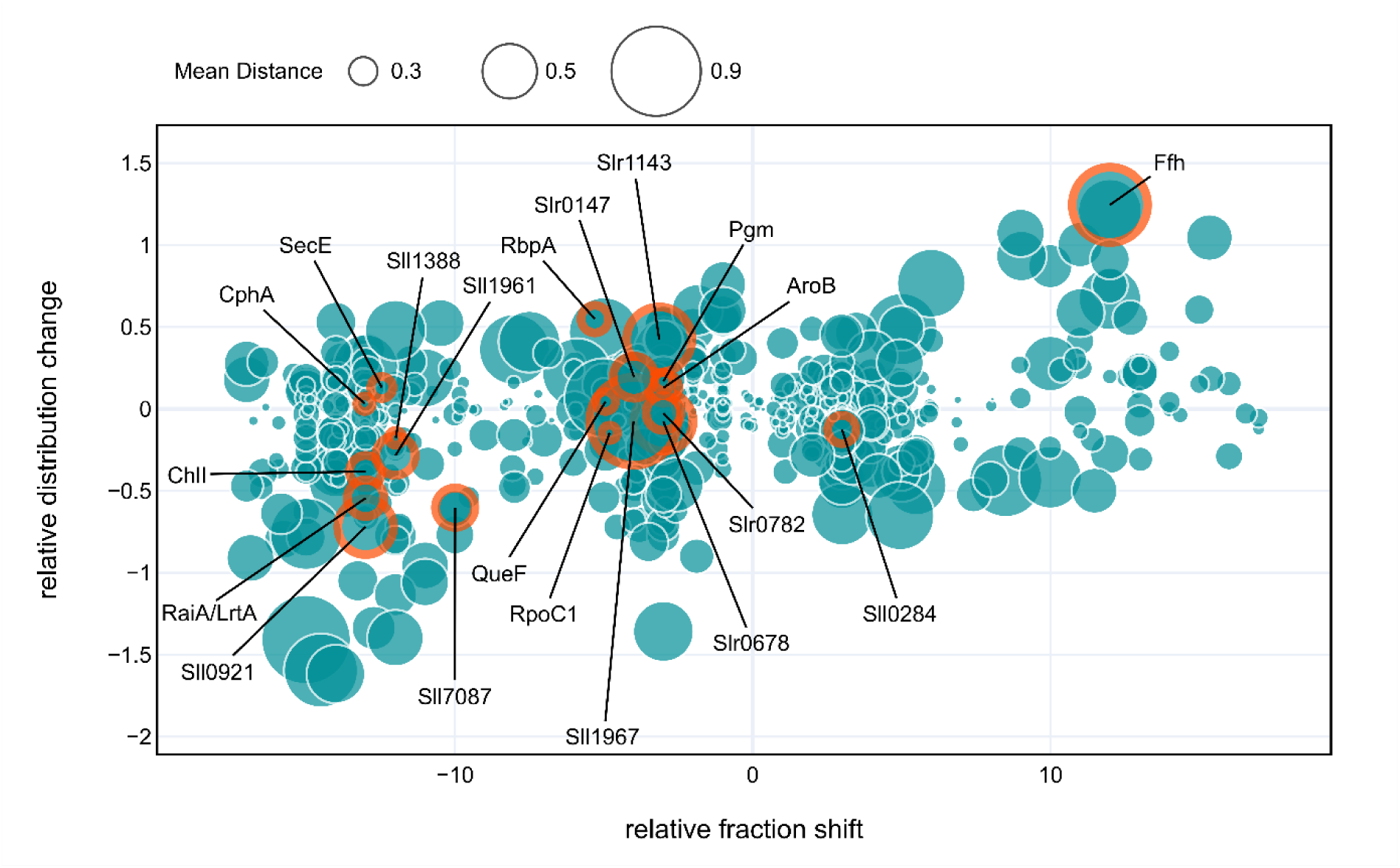
Visualization of shift types from *Synechocystis* Grad-R data. Each bubble represents a single protein identified by mass spectrometry, the bubble size corresponds to the mean JSD distance. The x-axis shows the relative fractional shift, while the y-axis shows the entropy gain (positive) or loss (negative) upon RNase treatment. This reflects whether the treatment led to a sharper or broader distribution of the protein (see Methods). Proteins that are promising candidates for RNA binding are circled in orange. Sll1967 is the 23S rRNA (uracil(1939)-C(5))-methyltransferase RlmD, Sll7087 is the type III-Bv CRISPR-Cas protein Cmr4^90^, Hpf is the ribosome-associated translation inhibitor RaiA/LrtA (Sll0947)^55^, Sll1388 is an universal stress protein, Slr1143 is a diguanylate synthase^65^, see **Table 1** for further details.

**FIG 6.**
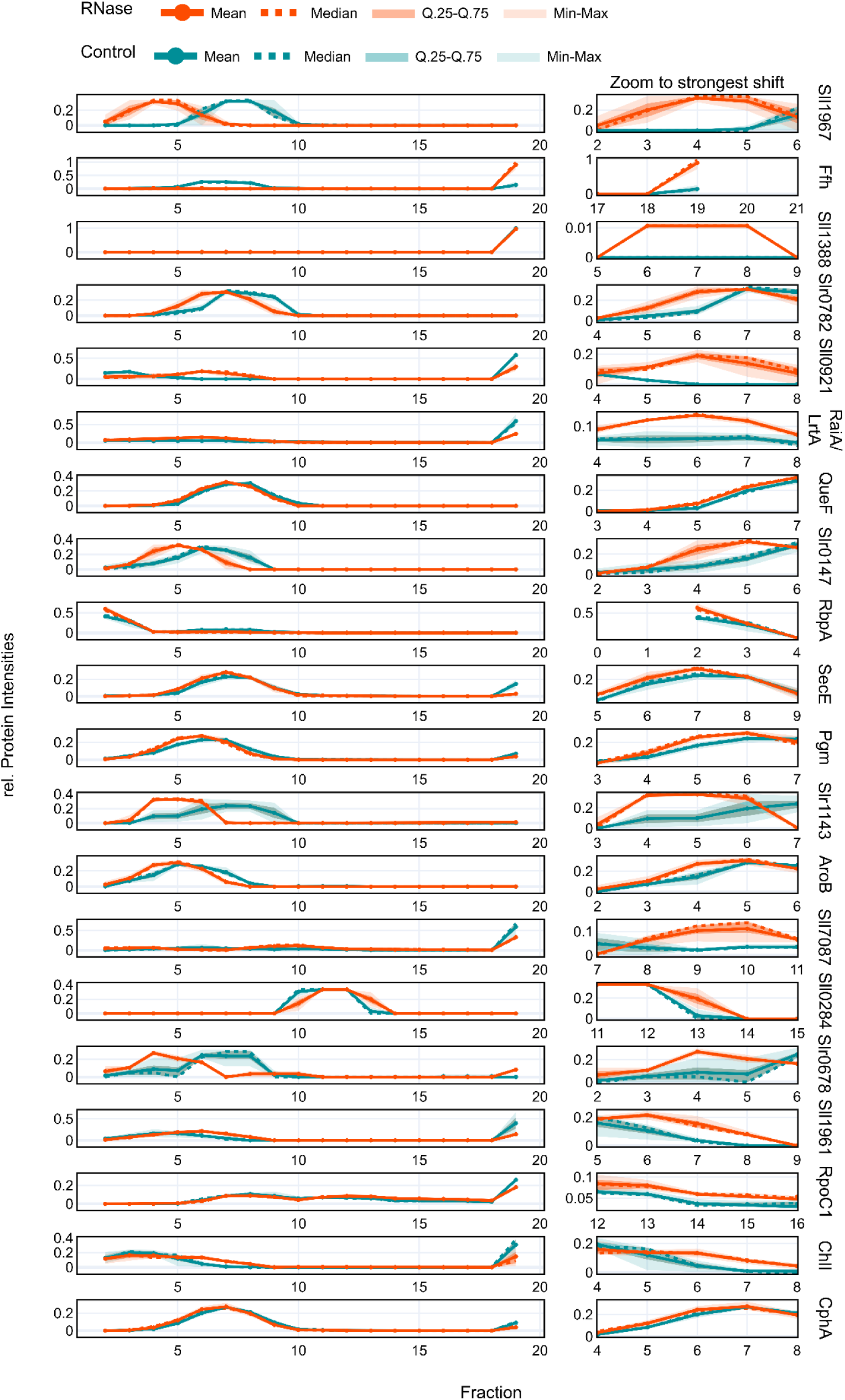
Distribution profiles of RNA-interacting protein candidates. Left panel: global view, right panel: zoom to the strongest shift upon treatment identified via the soft argmax of the position-wise relative entropy to the mixture distribution.

The distribution profiles of the selected candidates showed a variety of larger protein shifts, as well as smaller quantities of a respective protein shifting detected by RAPDOR **(Fig. 6)**. For instance, the RRM protein RbpA (Rbp1, Sll0517) showed only a small shift (**Fig. 6**), but RAPDOR annotated it with a high ANOSIM R value placing it on rank 33, showing that it has a good accuracy in detecting small shifts. This kind of small shift can also be seen in the distribution of Sll1388, a universal stress protein. Most of the protein was present in the pellet fraction, upon RNase digestion a small amount of the protein shifted to lower molecular weight fractions. However, the ANOSIM R value was 1, meaning that even though the amount of shifting protein was small, it was detectable in all replicates containing the protein. In addition to the ANOSIM R value, an RNA interaction was detected also for the homolog All1122 (42/61% identical and similar residues) in *Nostoc* 7120^47^. Moreover, an RNA chaperone function under cold-shock conditions was suggested for the homologous universal stress protein AtUSP (At3g53990; 24/44% identical and similar residues with Sll1388) in *Arabidopsis thaliana*^50^, highlighting the likely conserved interaction of these proteins with RNA. The comparison to the GradR dataset from *Nostoc* 7120 revealed with Slr2130 andSl0678 two further proteins on our short list which have homologs in *Nostoc* 7120 that showed RNA-dependent shifts^47^.

One of the most promising candidates with a distinctive shift **(Fig. 6)** was the TRAM domain containing protein Sll1967. It is annotated as a homolog of RmlD, which is involved in the methylation of tRNAs as well as rRNA in *Escherichia coli* and *Bacillus subtilis*^51–53^. However, experimental validation in cyanobacteria is lacking.

Another interesting candidate is the light-repressed transcript *lrtA,* initially described in *Synechococcus* sp. PCC 7002 as a rapidly induced gene when the cells were transferred from light to darkness^54^. Later, this observation was extended to include *Synechocystis* 6803^55^, where the association of the encoded protein Sll0947 with ribosomes was demonstrated as was its relevance in survival after stress exposure^56^. Structural modeling predicts this protein to be a homolog of the ribosome-associated translation inhibitor RaiA. Work in enterobacteria suggested RaiA (39/59% identical and similar residues with Sll0947) as inactivating 70S ribosomes and storing them as “sleeping ribosomes”^57^, consistent with its observed expression profile in cyanobacteria^54,55^.

Since it has become increasingly clear in recent years that especially enzymes involved in large metabolic pathways may possess moonlighting abilities to function as RBPs in addition to their enzymatic function^58–60^ we included three enzymes with a promising ANOSIM R value in the list of candidates. These proteins are part of the amino acid metabolism (Slr0782), the queuosine synthesis (QueF) and glycolysis (Pgm), which were placed by RAPDOR on ranks 22, 31 and 36 (**Table 1**).

In addition to RNA binding properties, potential DNA binding was also included in the analysis, as many bacterial DNA binding proteins can also bind RNA^61,62^, especially proteins harboring a helix-turn-helix motif such as the proposed candidates Sll0921 and Sll1961. With its involvement in photosystem 2 assembly^63^, Slr0147 is also of high interest as an RBP although it does feature a vinyl 4 reductase domain suggesting its involvement in chlorophyll metabolism. Some RBPs were shown to be involved in the transport of transcripts to the thylakoid membrane^29,64^. Therefore, it is possible that RBPs are involved also in the process of photosystem assembly. With its function as diguanylate cyclase the GGDEF domain containing Slr1143 protein possesses nucleotide binding capabilities^65^. The shift observed upon RNase treatment suggests that more than single nucleotides are bound to the protein. The GGDEF domain-containing CrsD is involved in the degradation of regulatory RNAs in *E. coli*^66^, while the RBP CrsA is involved in the tight regulation of diguanylate cyclases^67^. Consequently, an involvement of Slr1143 in RNA-regulated processes may be anticipated as well.

Another candidate of interest is the magnesium chelatase ATPase subunit I protein ChlI (RAPDOR rank 93, SVM score 112). Magnesium chelatase is a heterotrimeric enzyme (subunits ChlD, ChlI and ChlH) that performs the first committed step in the pathway to chlorophyll^68^. The catalytic subunit is ChlH^69^, which can also function as an anti-sigma factor^70^ showing its involvement in transcription-related processes. Our data suggest an RNA-binding capability for the subunit ChlI. Here performed structural alignments^71^ showed similarity to the eukaryotic DNA replication licensing factor MCM5, which was recently discovered as RBP^72^. Hence, an RNA-binding function of ChlI could well play a regulatory role, too.

An interesting case could be CphA, the enzyme that produces the nitrogen storage component cyanophycin and plays an important role in nitrogen metabolism. An association with RNA binding is given by the SVM score and the shift of the protein from the pellet fractions to the lower molecular fractions upon RNase digestion **(Fig. 6)**. This finding is consistent with the results of a recent study on the *Synechocystis* YlxR homolog. In this study CphA was the most highly enriched protein interaction partner in the protein pulldown of YlxR, suggesting a tight interaction either with YlxR or with the RNA interaction partner of YlxR, *rnpB*, the RNA component of RNase P.

### Application of RAPDOR to spatial proteomics

To showcase its potential for further applications, we used RAPDOR to re-analyze existing spatial proteomics data. In spatial proteomics, the amount of protein in different cellular compartments is measured. Martinez-Val et. al.^38^ investigated proteome dynamics upon Epidermal growth factor (EGF) stimulation in HeLa cells and compared it to unstimulated cells. The entire proteome was assessed across distinct cellular compartments at different time points after stimulation. Each compartment comprised two fractions: cytosolic, membrane-bound organelles, and the nucleus/nucleolus. Such a setup results in six categories instead of numerical fractions, which is also supported by RAPDOR, but results in slightly different plots. Notably, their methodology diverges from GradR as it does not rely on gradient density centrifugation for fractionation.

The analysis pipeline used by Martinez-Val et. al.^38^ calculates a mobility score per protein. This score reflects the total percentage of shifting protein. In addition, it performs two moderated t-tests at the fractions with the highest loss/gain in protein amount respectively. The resulting two p-values are then combined using Fisher’s method in order to identify proteins with a significantly different distribution after EGF treatment. Here we show that an analysis using RAPDOR yields similar results when calculating the *JSD* instead of a mobility score. Therefore, we used pre-analyzed data from Martinez-Val et al.^38^ as input for RAPDOR. This data was already filtered and imputed. Since the dataset included four replicates per condition, it was possible to calculate p-values using RAPDOR’s global ANOSIM mode (see Methods). However, since the power of the test is still weak for four replicates we decided for a less strict cutoff and used an adjusted p-value of 0.1, thus accepting a higher false discovery rate. Our results indicate that for each treatment timepoint most of the proteins identified in the original publication also showed a high *JSD* of at least 0.2 as well as a significant shift when analyzed via RAPDOR (**Fig. 7A**). Especially the shifts of GRB2, CBL, and SHC1 at two and eight minutes after EGF treatment were captured by RAPDOR. Those shifts indicate the recruitment of receptor tyrosine kinase (RTK) adaptor proteins as already highlighted^38^. Interestingly the analysis pipeline used in the original publication did not detect some clearly shifting proteins including FOXJ3. This showed a shift at all time points, which was most likely missed because the original publication based the statistical test on selecting the fractions with the highest loss or gain. In contrast, our approach does take all fractions into account by comparing distributions. Here, FOXJ3 accumulated in membrane-bound fractions and reduced its presence in the others (**Fig. 7B**). This leads to a low loss in the fraction with the largest loss and can result in a false negative observation when using t-tests to identify a substantial reduction over all replicates at this fraction. Although nothing is currently known about FOXJ3’s involvement in EGF signaling, RAPDOR has identified two proteins that are already known to be linked to this pathway. The endocytic scaffolding protein intersectin (ITSN1) and microphthalmia-associated transcription factor (MITF) both showed shifts from the nucleus towards cytosolic fractions at timepoints 2, 8, and 20 min that were not captured using the original analysis pipeline. While MITF is proposed to negatively regulate epidermal growth factor receptor (EGFR) expression^73^, ITSN1 is known to mediate EGFR signaling through modulating its ubiquitylation via the RTK adaptor CBL^74^. These findings underscore the enhanced sensitivity and accuracy of the RAPDOR method in detecting protein shifts, revealing novel insights overlooked by previous analyses.

**FIG 7.**
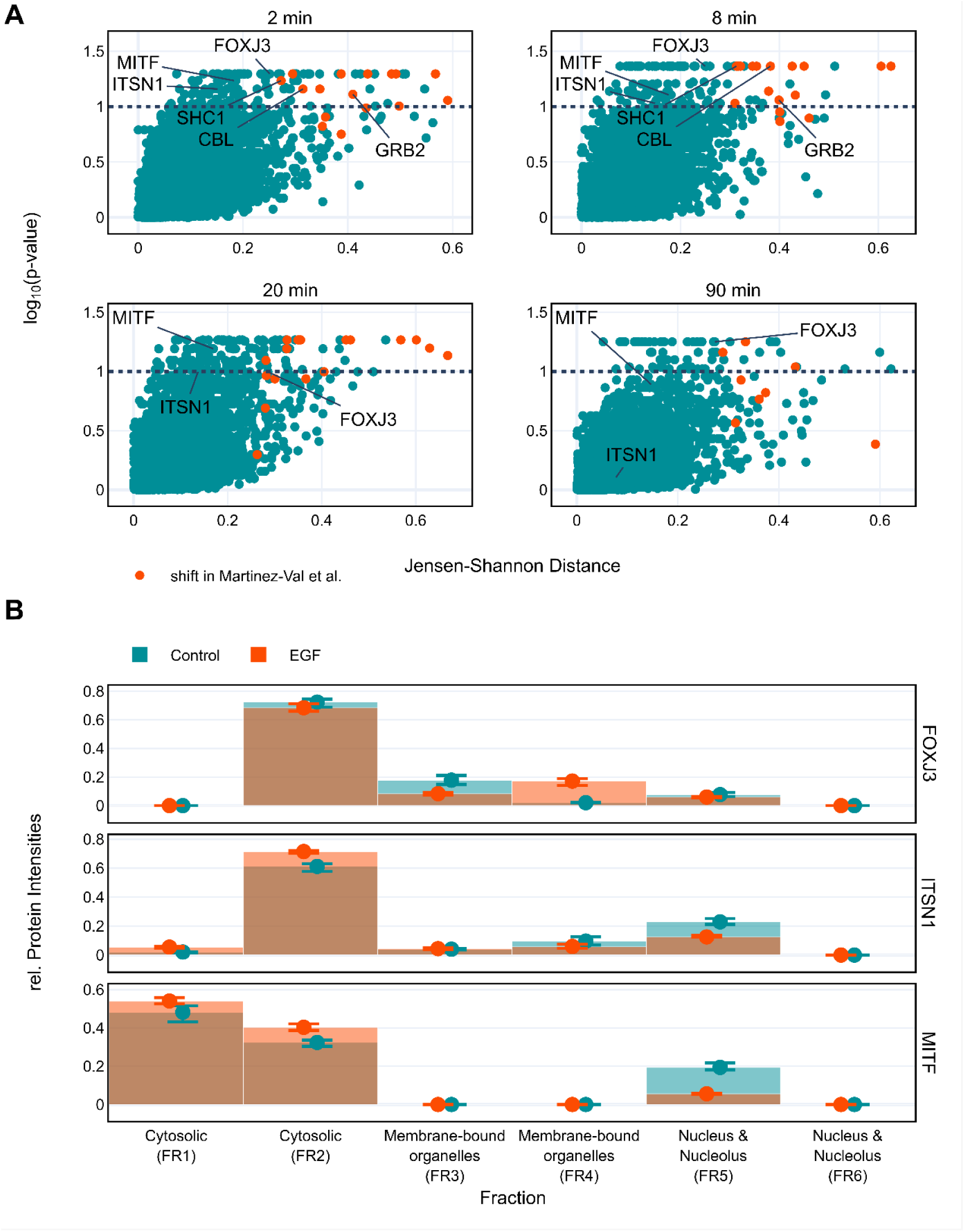
Analysis of spatial protein redistribution upon EGF treatment. (**A**) Protein shifts at 2, 8, 20, and 90 min of EGF treatment. Black dotted lines indicate typical cut-off values for such an experiment. Orange dots represent proteins that were identified with a distribution significantly different from the original analysis^38^. **(B)** Redistribution of FOXJ3, ITSN1 and MITF at 2 min of EGF treatment. After treatment, FOXJ3 showed a significant accumulation in membrane-bound organelles (FR4). In contrast, ITSN1 and MITF shifted towards cytosolic fractions. These shifts were not captured by the original analysis pipeline^38^.

Another problem might arise from the fact that combining two p-values using Fisher’s method, as done in the original publication, assumes their independence and is known to produce too low p-values for dependent tests. However, since the tests here are based on the same protein distribution, they are highly dependent. On the other hand, using the mobility score cutoff most likely counteracted this problem. In contrast, RAPDORs global ANOSIM only assumes that the R values of different proteins follow the same distribution. This seems to be valid for all four time point comparisons when observing the histograms of R values (**Supplementary Fig. S4**).

Wang et al.^39^ generated a dataset on the spatial redistribution of proteins for *Synechocystis* 6803. Their experiment focused on the re-distribution of proteins within cells upon different abiotic stress treatments. Since *Synechocystis* is a prokaryotic model organism and thus has no nucleus, their dataset only contains information about membrane-associated and cytosolic proteins. We created a JSON file compatible with RAPDOR for each comparison of the control treatment with one of the stress responses, which are nitrogen starvation, iron deficiency, cold, heat, and darkness. However, each comparison only had three replicates and consequently we used a ranking instead of p values. Noteworthy, main findings of the publication were reproduced using RAPDOR, such as the redistribution of some ribosomal proteins towards the membrane upon cold stress (**Supplementary Fig. S6)**.

These findings indicate the applicability of RAPDOR for experiments different than in-gradient profiles, and the analyzed datasets are promising starting points for other researchers to detect further protein redistributions. For visualization, all the supplementary JSON files can be plugged into local versions of RAPDOR, provide her as **Supplementary File 1**.

## Discussion

### RNase sensitive gradient fractionation of *Synechocystis* and peak-less shift identification

Gradient fractionation of RNase-treated samples to identify novel RBPs was previously performed with extracts from mammalian cells^27^ as well as from *Salmonella*^28^. Here we show that the method could successfully be applied to *Synechocystis*, a unicellular cyanobacterium. We verified the RNase-induced shift that is crucial for this method by analyzing ribosomal proteins of the small and large subunit (**Fig. 2**). The analysis of the entire dataset benefitted from the here developed RAPDOR workflow that allowed the peak-less shift identification.

The fact that R-DeeP detected fewer shifting ribosomal proteins than RAPDOR highlights an important drawback of the former compared to the latter approach. The majority of ribosomal proteins showed a change in the spread of the distribution rather than a shift. Even though this redistribution was reproducible among the replicates, the control and RNase-treated peaks were approximately at the same position. Such a behavior is not captured via the R-DeeP pipeline. In contrast, the probability-distance-based approach of RAPDOR can detect such shifts, mainly because it does not depend on peaks and their positions, but rather on changes in the distribution, thus taking all fractions into account. While R-DeeP assumes that proteins follow a multi-Gaussian distribution along the gradient, its authors are also aware of the fact that this might not be the case for all proteins. Therefore, some proteins become tagged to indicate a problematic fit. In addition, the R-Deep approach uses a t-test at each peak, which assumes that the normalized amount of protein follows a gaussian distribution. While a t-test is robust against violations of this assumption for large sample sizes, three replicates per group might produce misleading results when assumptions are not met. In contrast, RAPDOR’s non-parametric approach makes no assumptions about an underlying distribution, neither along the fraction dimension, nor at the peaks. This is also not the case when using the ranking approach which is helpful if the number of replicates is not sufficient to gain significant results just like in our dataset. Even though this ranking might not fully adjust for different variances of the distribution-data of different proteins, Clarke et. al. already pointed out that the R value itself can be used as a comparative measure^44^. Further, we show that the ranking might be adequate to identify RNA-dependent proteins as we found known RBPs within the first ranks of our dataset. Besides ribosomal proteins, these were RBPs such as the signal recognition particle protein Ffh on rank 17, the CRISPR protein Cmr4 on rank 51, or the RRM domain protein RbpA on rank 33. RbpA was ranked promisingly by RAPDOR despite the low mean distance of 0.26 between the treated and untreated samples because the shift was reproducible among the replicates resulting in a high ANOSIM R value. Cmr4 was demonstrated to bind crRNA and target RNA cleavage in type III-B CRISPR-Cas complexes^75^

We are aware of the fact that the RAPDOR workflow may also miss potential RNA-dependent proteins as false negatives, especially when using strict cutoff values for the ANOSIM R. This is illustrated here by Rps1A and Rps1B encoded by *slr1356* and *slr1984*, respectively. Rps1 is thought to participate in recruiting mRNA to the 30S subunit^76^, but homologs exist only in some bacteria. Rps1A and Rps1B in *Synechocystis* 6803 are involved in the Shine–Dalgarno-independent initiation of translation^77,78^ and therefore are known RNA-dependent proteins. Rps1b was captured by RAPDOR, on rank 53, while Rps1a was placed on rank 103, with an ANOSIM R value of 0.407, below our cutoff. On the other hand, missing information on the set of true-positive RBPs limits the ability to derive reasonable cutoff values and benchmark tool performance in an unbiased manner.

Despite some exciting examples of RBPs detected by RAPDOR, the total number of RBPs remains likely gravely underestimated. For example, the majority of proteins annotated with the RNA-binding GO term were not experimentally verified here. The reasons certainly include a lack of resolution. We estimate that the molecular mass difference required for a protein to shift by one fraction is at least 50 kDa (**Supplementary Fig. S2)**. Thus, only proteins binding to RNA at least 150 nt in length can shift by one fraction if this RNA is entirely degraded. Another important aspect may be that only a fraction of an RBP was actually bound to RNA, while a larger fraction was not. Finally, some proteins sharing the GO annotation simply had an insufficient number of replicates with a signal in the mass spec data. In contrast, the fact that TriPepSVM detected all the annotated RBPs likely indicated bias from training. GO-annotations in *Synechocystis* mainly stem from homology inference, which is also the kind of data the SVM model was trained on.

### The RAPDOR workflow is broadly applicable

One focus of our research was to develop a tool that is easily applicable for similar experiments. This has two aspects. First, we use a non-parametric approach, which does not make any assumption about the form of the distribution of the profiles and, to best of our knowledge, is the first application of ANOSIM in this field. In both GradR data and in the spatial proteomics example, the previous analysis was either based on a parametric modelling of the distribution itself or involved parametric tests like the student’s t-test on derived values. Second, implementational aspects are also important for a broad applicability. Hence, the RAPDOR pipeline is optimized for low runtime and memory consumption, making it suitable for average consumer laptops. Further, the tool is supported by comprehensive online documentation for the Python API and the Dash graphical user interface (GUI), featuring tutorials on setting up the tool with custom datasets. Such datasets can be analyzed either locally via Python or in the GUI directly (**Fig. 8**). It is then possible to set up a server to host such pre-analyzed data via a configuration and a JSON file. This disables the computationally more expensive analysis and provides a direct way to visualize data along with a publication. It also allows for more in-depth and visual analysis of individual distribution changes between different conditions. Consequently, we set up a webserver to display the *Synechocystis* GradR data under the following link: (https://synecho-rapdor.biologie.uni-freiburg.de)

**FIG 8.**
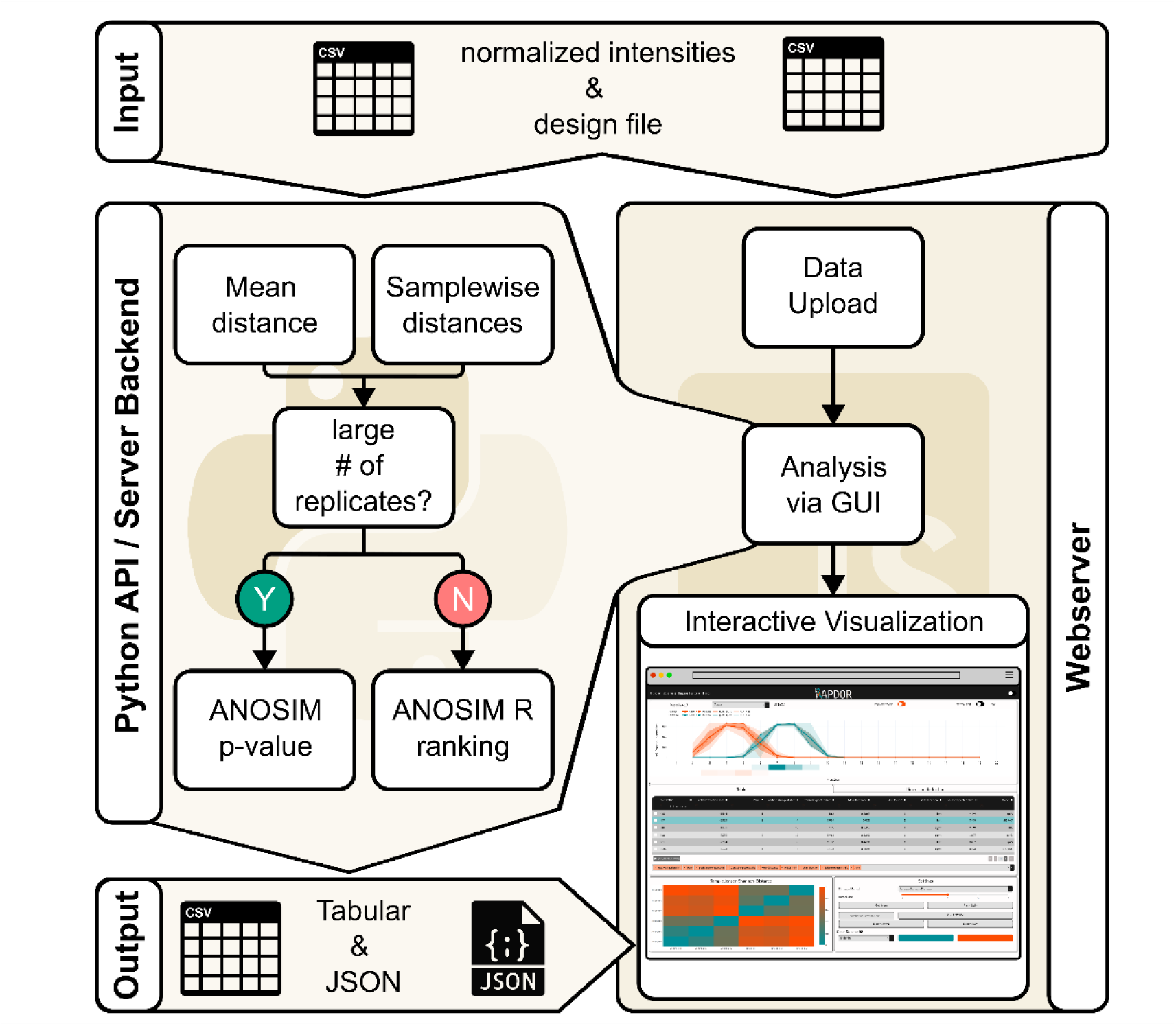
RAPDOR supports multiple usage methods. RAPDOR has the potential to conduct analysis either directly in python or as an interactive webserver. The python-analyzed data can be exported as a JSON file and displayed in the webserver environment afterwards.

To demonstrate the potential to distribute pre-analyzed data, we analyzed publicly available datasets, which focus on the re-distribution of proteins across cellular compartments of different model organisms. We offer these datasets in JSON format, compatible with the RAPDOR visualization tool, facilitating their utilization by other researchers for hypothesis generation (**Supplementary File 1**).

### Materials and Methods Culture conditions

Triplicate cultures of *Synechocystis* 6803 PCC-M^79^ were maintained in 100 mL BG11^80^ supplemented with 20 mM 2-{[1,3-Dihydroxy-2-(hydroxymethyl)propan-2-yl]amino}ethane-1-sulfonic acid (TES) pH 7.5 under continuous white light of 50 µmol photons m^−2^ s^−1^ at 30°C.

### Cell lysis and RNA removal

Cells were harvested after reaching an OD_750_ of 0.9 by centrifugation (4,000 *x* g, 4°C, 20 min). The cell pellet was resuspended in 800 µL lysis buffer + 1 mM DTT (20 mM Tris/HCl (pH 7.5), 150 mM KCl, 10 mM MgCl_2_) containing protease inhibitor (c0mplete EDTA-free protease inhibitor, Roche). Cell lysis was performed by mechanical disruption using a pre-chilled Precellys homogenizer (Bertin Technologies). To remove unlysed cells and glass beads, samples were centrifuged (500 *x* g, 2 min, 4°C), and the supernatants collected for further processing. To solubilize membrane proteins, cell lysates were incubated in the presence of 2% n-dodecyl β-D-maltoside for 1 h in the dark at 4°C. Cell debris was then removed by centrifugation (21,000 *x* g, 4°C, 1 h). The cleared lysate was divided into two fractions of equal volume. One of the fractions was incubated with 100 µL RNase A/T1 mix (Thermo Fisher Scientific) for 20 min at 22°C. The control fraction was treated with 100 µL mock buffer (50 mM Tris-HCl (pH 7.4), 50% (v/v) glycerol) + 0.4 U of RNase inhibitor (RiboLock, Thermo Fisher Scientific) for 20 min at 22°C. Samples were then kept on ice until loading.

### Gradient preparation and fractionation

Gradients were prepared using the Gradientmaster 108 (Biocomp) to obtain a linear gradient of solution 1 (10% (w/v) sucrose in lysis buffer) and solution 2 (40% (w/v) sucrose in lysis buffer). Open-Top Polyclear^TM^ Centrifuge Tubes 9/16 x 3-1/2 in. (Seton Scientific) were used as centrifugation tubes. Gradients were overlayed with 400 µL of solution 1 before loading 500 µL lysate. Separation of the lysate was achieved by ultracentrifugation using a swinging-bucket rotor (Beckman SW40 Ti) for 16 h at 285,000 x *g*. 20 fractions of equal volume (∼ 600 µL) were collected using the PGF ip Piston Gradient Fractionator (Biocomp), except for the pellet fraction (fraction 20), which was collected manually.

### Sample preparation and details of mass spectrometry measurements

One hundred µL of each fraction was used for mass spectrometric analysis of gradient samples. Samples were mixed with 4x sample volume of 50 mM triethylammonium bicarbonate (TEAB) and reduced by addition of 0.5 µM Tris(2-carboxyethyl)phosphine (TCEP) followed by incubation at 37°C for 45 min under constant shaking at 900 rpm. The samples were then alkylated with 0.5 µmol iodoacetamide (15 min, room temperature, dark). Proteolytic digest was achieved by adding 1 µg trypsin (Promega) and incubating the samples for 16 h at 37°C, 900 rpm. The digestion was stopped by adding 1/26 volume of 50% (v/v) trifluoroacetic acid. Peptides were purified after digestion using Pierce C18 Tips (Thermo Fisher Scientific) according to the manufacturer’s protocol. For fractions with the highest protein content (1-10 and 20) the purifying procedure was repeated twice, and the eluates were pooled. Recovered peptides were dried and resuspended in 20 µL of 0.1% acetic acid using an ultrasonic bath. To monitor reproducibility of LC-MS runs, retention time calibration peptides (iRT, Biognosys) were spiked in a 1:100 ratio.

LC-MS/MS analyses were performed on an LTQ Orbitrap Velos Pro (ThermoFisher Scientific, Waltham, MA, USA) using an EASY-nLC II liquid chromatography system. Tryptic peptides were subjected to liquid chromatography (LC) separation by loading them on a self-packed analytical column (OD 360 μm, ID 100 μm, length 20 cm) filled with 3 µm diameter C18 particles (Dr. Maisch, Ammerbuch-Entringen, Germany). Peptides were eluted by a binary nonlinear gradient of 2–99% acetonitrile in 0.1% acetic acid over 88 min with a flow rate of 300 nL/min and subsequently subjected to mass spectrometry (MS). For MS analysis, a full scan in the Orbitrap with a resolution of 30,000 was followed by collision-induced dissociation (CID) of the twenty most abundant precursor ions. MS2 experiments were acquired in the linear ion trap.

Database searches were performed against all proteins predicted to be encoded in the *Synechocystis* 6803 chromosome (NC_000911.1) and the four plasmids pSYSA, pSYSG, pSYSM, and pSYSX (NC_005230.1, NC_005231.1, NC_005229.1, and NC_005232.1, respectively). The database was supplemented with sequences of known sORFs and RNase A/T resulting in a total of 3743 entries. Database search as well as label-free protein quantification was performed using MaxQuant (version 2.0.3.0)^81^. Common laboratory contaminants and reversed sequences were included by MaxQuant. Search parameters were set as follows: trypsin/P specific digestion with up to two missed cleavages, methionine oxidation and N-terminal acetylation as variable modification, carbamidomethylation at cysteines as fixed medication, match between runs with default parameters enabled. The FDRs (false discovery rates) of protein and PSM (peptide spectrum match) levels were set to 0.01. Two identified unique peptides were required for protein identification. LFQ^82^ and iBAQ^83^ were exported as quantitative values of protein abundance.

The generated MS data have been deposited to the ProteomeXchange Consortium via the PRIDE partner repository with the dataset identifier PXD045848.

Polyacrylamide gel electrophoresis and Western blotting 20 µL of each fraction was boiled with 1x protein loading buffer (Tris/HCl 0.5 M (pH 6.8) 50 mM, SDS 2%, glycerol (v/v) 6%, DTT 2 mM, Bromophenol Blue (w/v) 0.01%) at 95 °C for 10 min and then separated by 15% SDS-polyacrylamide gel electrophoresis (SDS-PAGE)^84^.

### RNA isolation and Northern blotting

RNA extraction and northern blotting were performed as described previously^25^. Primer and oligonucleotide sequences are listed in **Supplementary Table S6**. Signals were visualized using Typhoon FLA 9500 (GE Healthcare) and Quantity One software (Bio-Rad).

### Jensen-Shannon Distance

The Jensen-Shannon Distance (*JSD*, see Eq. 3(3)) is the default effect size measure for an individual replicate of the RAPDOR tool. For two probability distributions P and *Q*, the *JSD* is defined as follows:

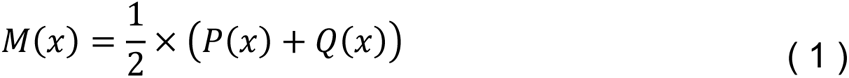

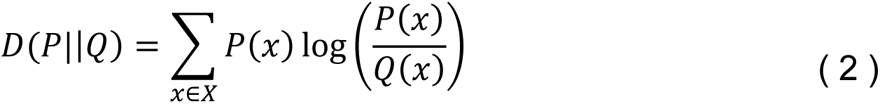

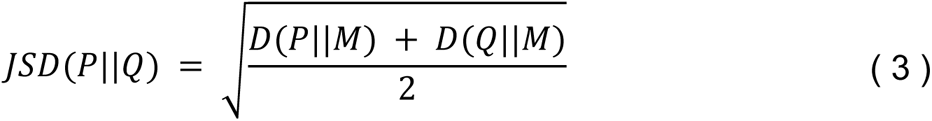

Here, *M* refers to the mixture distribution of P,*Q* and *D* is known as the Kullback-Leibler (KL) divergence (see Eq. 1 and 2(3)), which is a commonly used measurement on probability distribution. In contrast to the more prominent KL divergence, which is not symmetric, the *JSD* is a metric function. This makes the *JSD* suitable for the downstream analysis for a non-parametric test of similarities between conditions (see the section on the ANOSIM R value below).

### Expected position of the strongest shift

For each treatment *t* the position of the strongest shift *S*_*t*_ is determined using the position wise relative entropy, which has its origin in convex programming (Eq. 4(4)). We used it since its sum is equal to the KL divergence, given that both P and *M* are probability functions^85^. In more detail, it is based on likelihood ratios of either treatment and the mixture distribution at position *x*, weighted by the probability P(*x*) of observing fraction *x*. To determine the position of strongest shift, we use a localization operator (Eq. 5) that corresponds to the temperature-scaled soft-argmax function, which is often used in machine learning. It counteracts uncertainty in our data and performed better than the argmax function for distributions with broader peaks. Basically, the soft-argmax function returns the expected position of the strongest shift. β is the hyperparameter for temperature scaling and must be carefully chosen for a given dataset.

The relative fraction shift (Eq. 6) is then obtained via a subtraction of the expected position of the strongest shift of the control samples (*t* = −) from the RNase treated one (*t* = +).

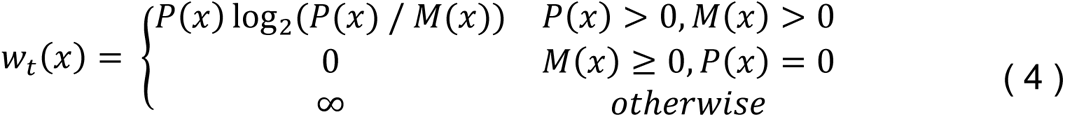

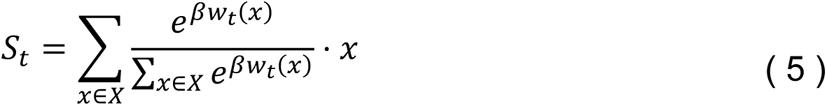

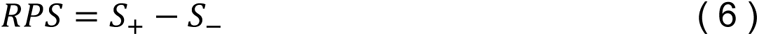

### Non-parametric test analysis of similarities (ANOSIM)

The non-parametric test analysis of similarities (ANOSIM) calculates a test statistic called ANOSIM R via a ranked distance matrix^44^. This is achieved via labeling fields in this matrix with two distinct labels depending on whether the distance originated from two samples of the same treatment *W* or a different treatment *B*. The R value for *n* samples is then calculated via the following equation:

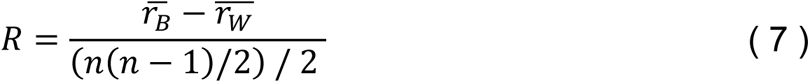

Hereby 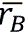 is the average rank of the different treated fields (e.g. in GradR control vs RNase) and 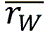 refers to the average rank of fields from the same treatment (e.g. in Grad RNase vs. RNase or control vs control). Via shuffling of the labels (*B*, *W*) it is possible to estimate a distribution over the R values and thus calculate a p-value.

While ANOSIM is a well-known statistical test for community ecologists, the procedure can in principle be adapted to any kind of multivariate data analysis, as long as there is a sufficient number of replicates per condition. However, three replicates are not sufficient, to achieve a p-value below a significance level α ≤ 0.05. This is because the number of permutations given a balanced sample layout with two conditions follows Eq. 8(8) thus resulting in only 10 distinct permutations for *n* = 3.

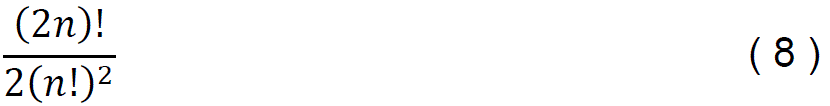

Depending on the number of replicates and instances (here proteins), it is possible to calculate p-values using the R value distribution of all instances as background. This method, further termed global mode, assumes that R values of different instances follow the same distribution. While this may not always be the case, such assumptions are generally robust and are employed by other tools. For example, the differential expression tool limma utilizes information from all genes for its underlying moderated t-test^86^. Further, the ANOSIM test statistic is bound between [−1,1]. This makes it comparable between proteins with the same number of samples, while it is not influenced by the effect size due to its non-parametric nature. Values close to 1 show that the distances between the same treatment are lower than the distances between different treatments. In contrast, –1 shows the opposite, and values around zero indicate evenly distributed distances. This offers the possibility to rank proteins according to a decreasing R value to identify those with reproducible shifts.

### Runtime & Memory consumption benchmarking

The runtime and memory consumption analysis was performed on a single Intel i5-10210U CPU core. Additionally, a dataset with more replicates was simulated by duplicating the existing measurements, resulting in a dataset with nine replicates each. This was used exclusively for runtime and memory consumption benchmarking.

### Support vector machine classification of candidate RBPs

In addition to the experiment-based approaches, we used TriPepSVM^19^, to predict candidate RBPs. TriPepSVM relies on the idea that a support vector machine (SVM) can make a prediction on whether a protein is an RBP solely based on amino acid sequence. The basic principle is creating positive and negative datasets, which will then train the SVM, leading it to predict RBPs based on sequence alone. A protein sequence is cut-up into overlapping k-mers, which form a vector. This vector is analyzed by the learning algorithm, and tripeptides are scrutinized by occurrence within the protein. This leads the SVM to make a prediction based on likelihood of said protein belonging to the RBP or non-RBP category. We have trained TriPepSVM for cyanobacterial genomes and applied it to two use cases, on *Nostoc* sp. PCC 7120^47^ and on *Synechocystis* 6803 (this work). To train the algorithm for cyanobacterial RBP candidates, the predicted set of proteins encoded in 10 cyanobacteria, *E. coli* K12, and *Salmonella typhimurium* LT2 was downloaded from the UNIPROT database^87^. To avoid duplicate annotations or paralogs, CD-Hit^88^ was utilized to create a unique dataset and remove redundant proteins with a similarity >90%^88^. This was done for the 12 considered organisms yielding 1,151 unique RBP candidates that were merged into a positive dataset.

From the pool of remaining proteins, all proteins with an RNA-binding domain in the Pfam database or with annotation keywords related to nucleic acid binding in UNIPROT, or GO terms in QuickGO, were discarded. After filtering by CD-Hit a negative dataset of 33,860 unique non-RBP was obtained. The kmerPrediction.r script of TriPepSVM^19^ was modified for the implementation of the SVM by changing the package used in conjunction with “KeBABs”^89^ from “e1071” to “LiblineaR”. For the selection of the best combination of parameters, each dataset was randomly split into training (90%) and testing (10%) samples and used in a 10-fold cross validation by randomly sampling the subsets. The parameter combination resulting in the largest average balanced accuracy (BACC) was selected. We used a positive class weight of 2.7, a negative class weight of 0.05, cost = 1 and k-mer = 3. For further details see^47^. The proteins encoded by all 3.681 annotated protein-coding genes in *Synechocystis* 6803 were scored and a conservative threshold of 0.25 was selected as the SVM score for the classification as potential RBP. This prediction yielded a list of 306 candidate RBPs in *Synechocystis* 6803 (**Supplementary Table S3**).

## Supporting information

Supplementary Dataset S1 archive file

## Acknowledgments

We appreciate the support by the Deutsche Forschungsgemeinschaft (DFG, German Research Foundation) through the research training group “BioInMe” 322977937/GRK2344 to LH, DR, RB and WRH, DFG priority program SPP2002 “Small Proteins in Prokaryotes, an Unexplored World” (grants HE 2544/12-2 to WRH, BA 2168/21-2 to RB, and BE 3869/5-2 to DöB) and grant HE 2544/22-1 to W.R.H. MBA was granted by an Alexander von Humboldt postdoctoral fellowship. This study was supported by the German Research Foundation (DFG) under Germany’s Excellence Strategy (CIBSS – EXC-2189 – Project ID 390939984). We thank the IT department of Bio 3 for their help in setting up the web server.

## Authors’ contributions

WRH and LH designed the research. The experiments were performed by LH and VR. SM, PG, JB and DB carried out the mass spectrometry and related analyses. HR and MBA applied the TripepSVM approach and DR and RB developed the RAPDOR tool. All authors analyzed the data. WRH, LH, RB and DR wrote the manuscript with input from all authors. All authors read and approved the final manuscript.

## Data availability

The datasets produced in this study are available in the following databases:

● Mass spectrometry raw data were deposited at the ProteomeXchange (15) Consortium (http://proteomecentral.proteomexchange.org) via the PRIDE partner repository (16) under the identifier PXD045848.
● *Synechocystis* 6803 data accessibility and visualization: https://synecho-rapdor.biologie.uni-freiburg.de
● The RAPDOR tool is available as a pypi package and its documentation is hosted on https://domonik.github.io/RAPDOR/
● The code used to analyze the data including the modified R-DeeP script is available as a snakemake workflow on Github: (https://github.com/domonik/synRDPMSpec)

