## Supplementary Material for "RAPDOR: Using Jensen-Shannon Distance for the computational analysis of complex proteomics datasets"

[#]Co-sharing first authors

**Supplementary Information**

| | |
|---|---|
| **Supplementary Files:** | **p. 2** |
| **Supplementary Tables:** | **p. 2** |
| **Supplementary Figures:** | **p. 4** |
| **Supplementary References:** | **p. 10** |

### Supplementary Files

**Supplementary File 1.** This archive contains supplementary RAPDOR files used in the publication. All JSON files can be plugged into the RAPDOR Dash tool. Files are at least compatible with RAPDOR version 0.1.4. While they might work with other versions we cannot guarantee that.

To learn about how to install RAPDOR and upload those files visit:

https://domonik.github.io/RAPDOR/

### Files

#### synechoRAPDORGradRFile.json

The file for the *Synechocystis* 6803 GradR data. This is the same file that is displayed in our publication’s webserver.

#### HeLaEGFTreatment_egf_{x mins}.json

Pre-analyzed data for the reanalysis of data from Martinez-Val et al. (2021)[1]. Each file compares the protein distribution of HeLa cells x mins after EGF treatment to untreated cells.

#### RAPDORforSynechocystisabioticStress{stress}.json

The reanalyzed files for the publication Wang et al. (2023)[2] showing changes in distribution of membrane-bound vs. cytosolic proteins after different abiotic stress treatments. Stress conditions were cold, heat, darkness, nitrogen starvation (N) and iron starvation (F)

-- see separate archive file –

### Supplementary Tables

**Table S1. List of *Synechocystis* 6803 proteins detected by mass spectrometry.**

-- see separate Excel file --

**Table S2. Overview on GradR fraction complexity and distribution.**

Visualization of peak distribution within the biological replicates 1–3 (BR1–BR3) with or without (control) RNase. For each fraction, the number of proteins with the highest abundance in that fraction is shown. Orange indicates many peaking proteins, while turquoise indicates fewer peaking proteins.

-- see separate Excel file --

**Table S3. Classification as candidate RBP or not in *Synechocystis* 6803 according to the support vector machine classification.**

-- see separate Excel file --

**Table S4. Proteins with a high ANOSIM *R* value or an SVM prediction as RBP.**

-- see separate Excel file --

**Table S5. List of proteins redistributed upon cold stress after analysis by RAPDOR.**

-- see separate Excel file –

**Table S6. Deoxynucleotide primers used for the amplification of probe templates to detect sRNAs by Northern hybridization.** The name, ID and transcriptional unit (TU) of the respective sRNA are given as previously defined[3]. All sequences are given in 5' to 3' direction, the added T7 promoter sequences are indicated by lowercase letters.

| **Name** | **ID** | **TU** | **Fractions in Grad-seq** | **primer** | **Sequence** |
|---|---|---|---|---|---|
| PmgR1 | ncr0700 | 1715 | F5 – F11 | ncr0700_probe_fwd | CTTCACTGGCAGATAAAAAC |
| | | | | ncr0700_probe_T7_rev | taatacgactcactatagggGATGAACTGAGAAACAAACG |
| tmRNA/ ssrA | ncl1680 | 3496 | F6 - F11 | ncl1680_probe_fwd | AATGGTTTCGACAGGTTGGC |
| | | | | ncl1680_probe_T7_rev | taatacgactcactatagggCCCGTTTGAAGCTGACGATG |
| RNase P RNA/rnpB | 6803s01 | 139 | F8 – F12 | rnpB_probe_fwd | AGTTGCGGATTCCTGTCACAG |
| | | | | rnpB_probe_T7_rev | taatacgactcactatagggGTGGCACTGTCCTCACGCTC |

### Supplementary Figures

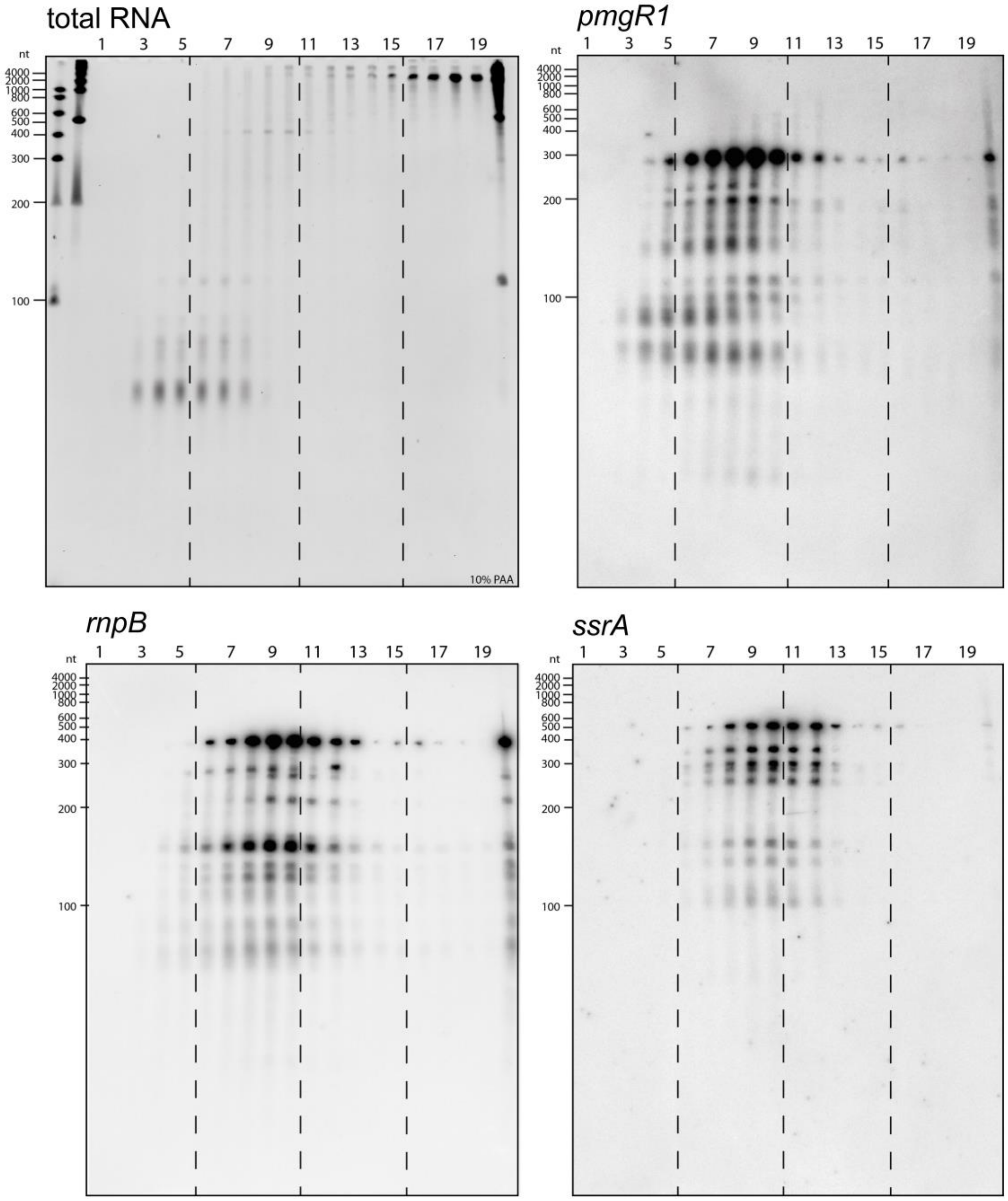


**FIG S1 Distribution of known RNA species in the untreated control samples.** Northern blots of 10% denaturing PAA gels hybridized with radioactively labelled probes. Gel image of total RNA prepared from the different gradient fractions as numbered on top. The molecular mass standards in the left two lanes were the High and Low Range RiboRuler RNA Ladders (Thermo Fisher Scientific). Hybridization is shown against the sRNA PmgR1[4], the RNase P RNA RnpB[5], and the transfer-messenger RNA (tmRNA) SsrA[6].

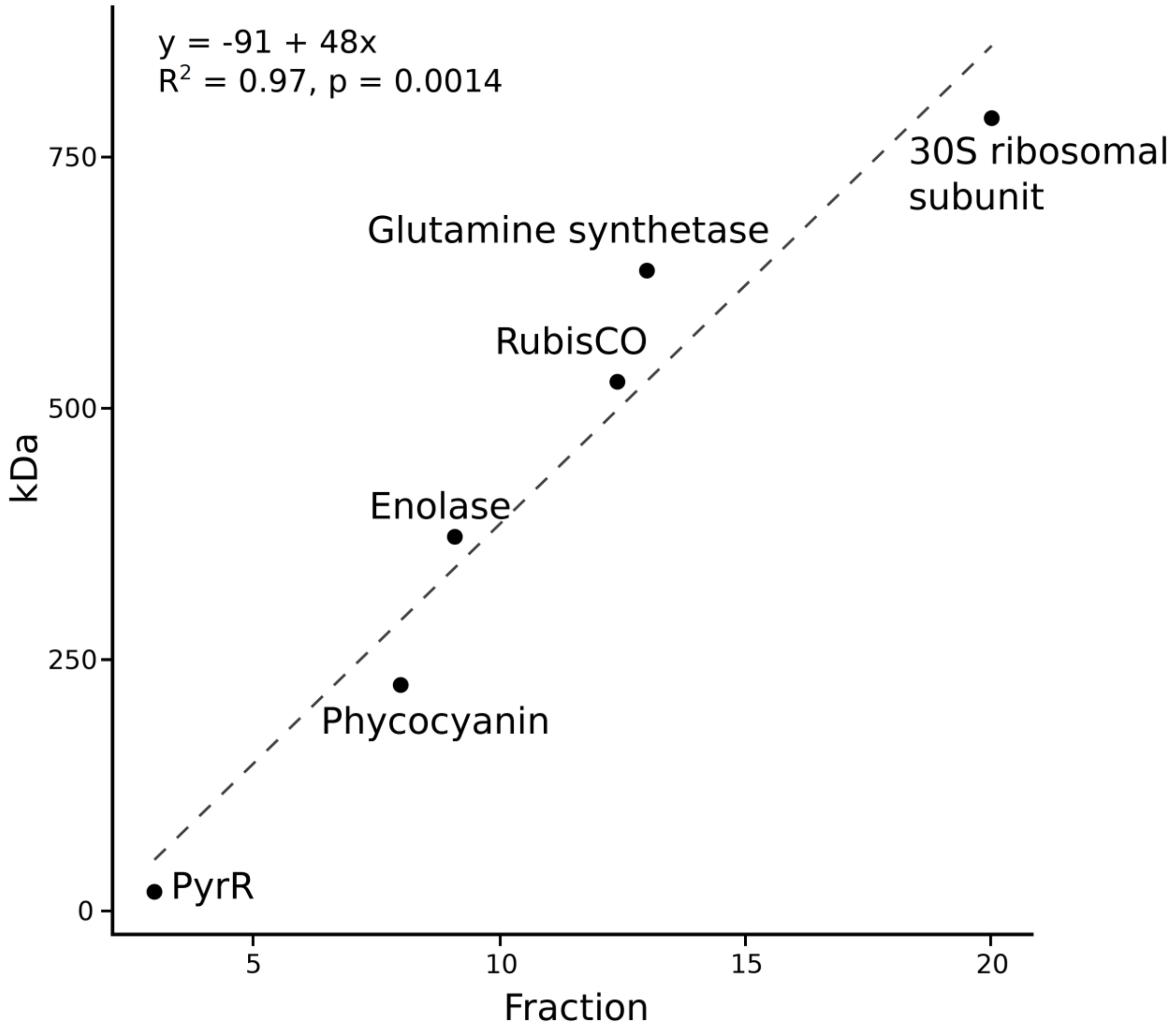


**FIG S2 Resolution of gradients.** Sedimentation of selected proteins and protein complexes in specific fractions of a sucrose density gradient after ultracentrifugation (x-axis) in comparison to the calculated molecular mass (y-axis). For the calibration curve, the respective peak fractions of the indicated proteins were selected. Masses were calculated for an α3β3 hexameric phycocyanin complex[7], for RubisCO consisting of an 8 small and 8 large subunits[8], for the homo 8-mer enolase *of Synechococcus elongatus*[9], the homo 12-mer glutamine synthetase[10] and 30S ribosomal subunit of *Escherichia coli*[11]. PyrR (Sll0368) is the bifunctional *pyr* operon transcriptional regulator/uracil phosphoribosyltransferase PyrR (molecular mass 19.946 kDa).

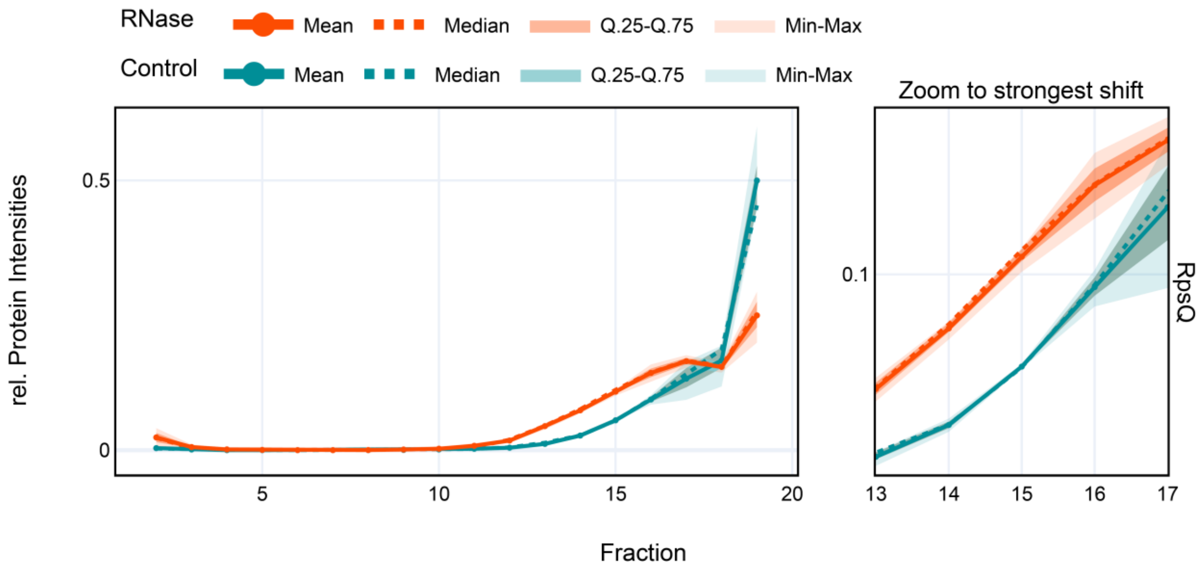


**FIG S3 Redistribution of RpsQ.** Like most small ribosomal subunit proteins, RpsQ showed a redistribution from the last fraction to nearby lower molecular mass fractions. This resulted in a broader peak that was not identified as a significant shift by the R-DeeP analysis pipeline.

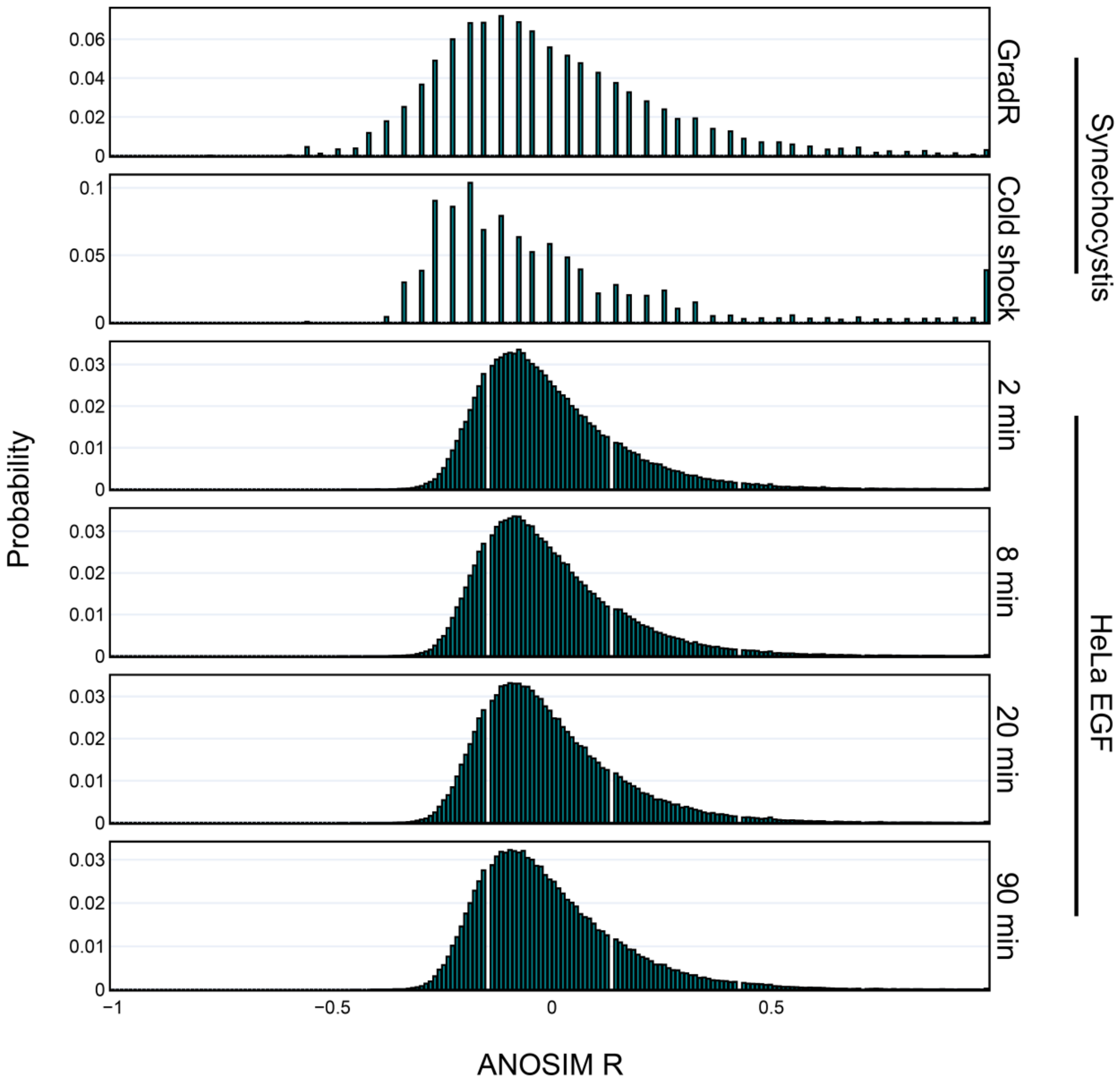


**FIG S4 Distribution of ANOSIM *R* values of the different datasets**. Distributions were generated using all possible permutation of treatment labels and calculating the ANOSIM *R* value. Afterwards, the values of different proteins with signal in all replicates were stacked together to generate the histograms of the background distributions. This method assumes that *R* values for different proteins follow a similar distribution. The two datasets from *Synechocystis* are the GradR dataset from this publication as well as the proteome reorganization from Wang et al.[2]. The EGF treated HeLa datasets correspond to different treatment time points from Martinez-Val et al.[1].

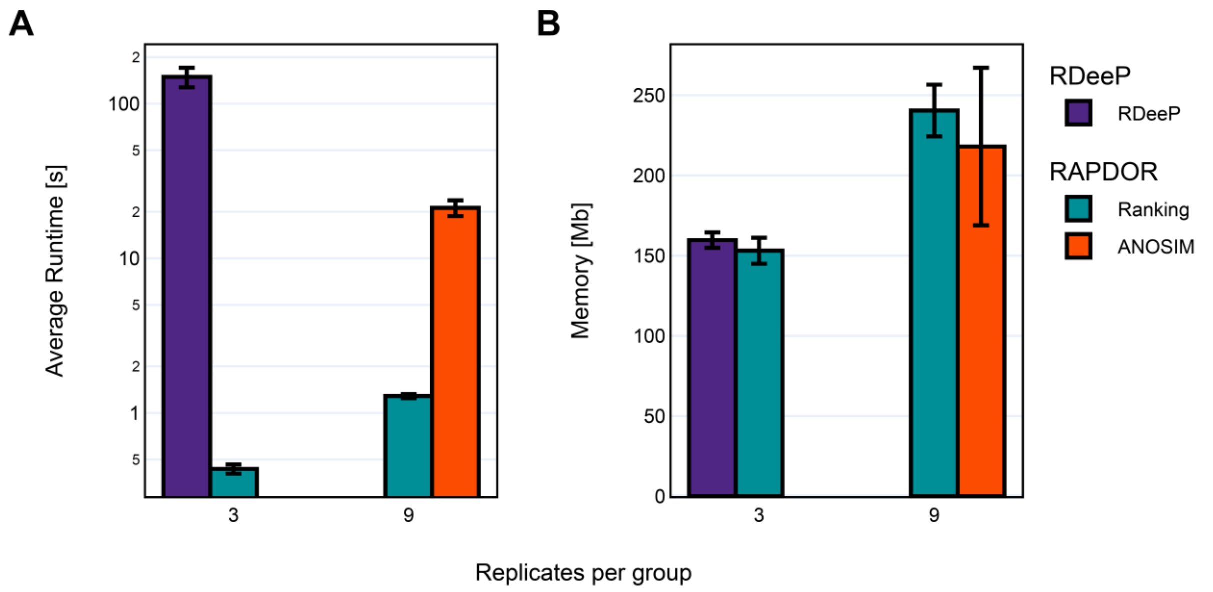


**FIG S5 Run time comparison of the tools. (A)** Average run time of 10 runs of each tool with different numbers of replicates. Note that the y-axis is on log scale. **(B)** Average memory consumption.

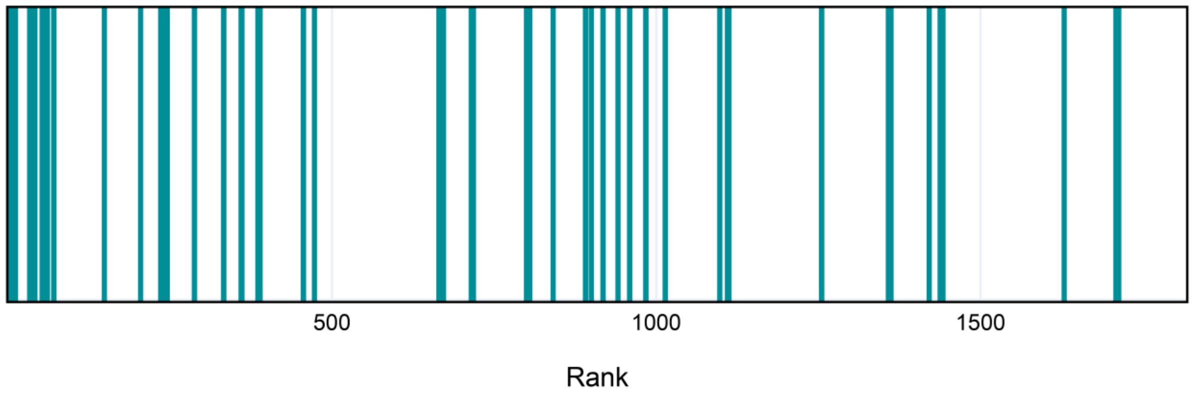


**FIG S6 Ranks regarding redistribution of ribosomal proteins upon cold stress from Wang et al.[2].** Ranks of ribosomal proteins that showed a shift towards the membrane fraction. Ranks are based on sorting the list of all proteins using a decreasing ANOSIM $R$ and a decreasing Jensen-Shannon-distance. Some ribosomal subunits showed a strong shift towards membrane upon treatment. The complete list or reanalyzed data on the redistribution of proteins upon cold stress is in **Table S5**.
